# The prospect of universal coronavirus immunity: a characterization of reciprocal and non-reciprocal T cell responses against SARS-CoV2 and common human coronaviruses

**DOI:** 10.1101/2023.01.03.519511

**Authors:** Mithil Soni, Edoardo Migliori, Jianing Fu, Amer Assal, Hei Ton Chan, Jian Pan, Prabesh Khatiwada, Rodica Ciubotariu, Michael S. May, Marcus Pereira, Valeria De Giorgi, Megan Sykes, Markus Y Mapara, Pawel Muranski

## Abstract

T cell immunity plays a central role in clinical outcomes of Coronavirus Infectious Disease 2019 (COVID-19). Therefore, T cell-focused vaccination or cellular immunotherapy might provide enhanced protection for immunocompromised patients. Pre-existing T cell memory recognizing SARS-CoV2 antigens antedating COVID-19 infection or vaccination, may have developed as an imprint of prior infections with endemic non-SARS human coronaviruses (hCoVs) OC43, HKU1, 229E, NL63, pathogens of “common cold”. In turn, SARS-CoV2-primed T cells may recognize emerging variants or other hCoV viruses and modulate the course of subsequent hCoV infections. Cross-immunity between hCoVs and SARS-CoV2 has not been well characterized. Here, we systematically investigated T cell responses against the immunodominant SARS-CoV2 spike, nucleocapsid and membrane proteins and corresponding antigens from α- and β-hCoVs among vaccinated, convalescent, and unexposed subjects. Broad T cell immunity against all tested SARS-CoV2 antigens emerged in COVID-19 survivors. In convalescent and in vaccinated individuals, SARS-CoV2 spike-specific T cells reliably recognized most SARS-CoV2 variants, however cross-reactivity against the *omicron* variant was reduced by approximately 50%. Responses against spike, nucleocapsid and membrane antigens from endemic hCoVs were more extensive in COVID-19 survivors than in unexposed subjects and displayed cross-reactivity between α- and β-hCoVs. In some, non-SARS hCoV-specific T cells demonstrated a prominent non-reciprocal cross-reactivity with SARS-CoV2 antigens, whereas a distinct anti-SARS-CoV2 immunological repertoire emerged post-COVID-19, with relatively limited cross-recognition of non-SARS hCoVs. Based on this cross-reactivity pattern, we established a strategy for *in-vitro* expansion of universal anti-hCoV T cells for adoptive immunotherapy. Overall, these results have implications for the future design of universal vaccines and cell-based immune therapies against SARS- and non-SARS-CoVs.

## Introduction

Coronavirus Infectious Disease 2019 (COVID-19) caused by Severe Acute Respiratory Syndrome Coronavirus 2 (SARS-CoV2), has resulted in over 6.5 million deaths worldwide[1]. SARS-CoV2 represents the third occurrence of a novel β coronavirus (CoV)-related disease in the last two decades. In 2003, SARS-CoV1 was identified in Asia, followed by Middle Eastern Respiratory Syndrome (MERS) CoV infection in 2012[2, 3]. These diseases had mortality rates of 9% and 40% respectively but have been confined to limited outbreaks [3-6].

Four endemic non-SARS human CoVs (hCoVs) circulate widely in the general population, including α-CoVs (hCoV 229E and NL63) and β-CoVs (hCoVOC43 and HKU1)[7, 8]. Endemic hCoVs cause up to 41% of seasonal upper respiratory infections[9, 10]. While respiratory illnesses caused by endemic hCoV are typically self-limited, hCoV infection can be severe and protracted in patients with co-morbidities such as stem cell transplant (SCT) and solid organ transplant (SOT) recipients, leading to hospitalizations, oxygen use, intensive care admissions and even death[11]. Thus, understanding endemic hCoVs infections is important, particularly for immunocompromised patients.

The role of cellular responses against SARS-CoV2 and other hCoVs is not fully understood[12], although emerging data suggest that robust T cell immunity correlates with rapid resolution of COVID-19[13-16]. Competent adaptive cellular immunity alone may be sufficient for eradication of SARS-CoV2 and recovery from COVID-19 in subjects with profound acquired or inborn defects in B cell function[17]. Patients with suppressed humoral immunity due to leukemia or lymphoma can mount an effective immune response to SARS-CoV2 if the T cell compartment is preserved[18]. Moreover, while vaccine or infection-induced neutralizing antibody titers have limited half-life, anti-CoV T cell memory might be long-lived[19], as documented in survivors of SARS-CoV1 infection who retained T cell reactivity over 10 years after recovery[20, 21].

Emerging data suggest that previously acquired heterologous memory responses might explain the very broad spectrum of COVID-19 manifestations and disease severity among infected subjects, as previous “common cold” induced immunity likely conveys at least partial protection[22, 23]. Unexposed individuals sometimes display SARS-CoV2-specific T cell reactivity, conceivably induced by previous exposures to hCoVs that share some common epitopes, [13, 24].

Recent work by Kundu et. al indicates that pre-existing T cell responses may correlate with resolution of COVID-19 without seroconversion [25]. However, the knowledge of T cell responses against the non-SARS hCoVs antigens remains limited, especially beyond the reactivity against S1 and S2 proteins targeted by vaccines. T cell reactivity against M and NP antigens from non-SARS hCoVs has not been fully explored.

Here we performed an in-depth analysis of CoV-specific T cell responses in healthy volunteers, (mostly healthcare workers, HCWs) with and without documented COVID-19 exposure as well as a cohort of high-risk immunocompromised patients (IP) including subjects with history of hematological malignancies, autologous and allogeneic SCT, SOT recipients and immunosuppressed patients with autoimmune diseases. T cell reactivity against the immunodominant antigens spike 1 and 2(S1, S2), membrane (M) and nucleocapsid (NP) proteins from SARS-CoV2 was characterized in relationship to responses against counterpart antigens from hCoV 229E, NL63, OC43 and HKU1. We also investigated cross-recognition of T cell responses against disparate epitopes derived from the spike antigen of multiple SARS-CoV2 variants detected during the pandemic[26], including the highly mutated omicron variant [27, 28]. Based on the observed cross-reactivity patterns, we postulate that a previously acquired infection or vaccination may provide broad T cell memory capable of recognizing, at least partially, future variants as well as related hCoV viruses. Finally, we established a strategy for *ex vivo* generation of universal multi-hCoV-specific T cells with enhanced ability to target common and emerging CoV[29]. We postulate that this T cell product may be useful in the clinic as adoptive transfer therapy or prophylaxis for CoV infections in SCT recipients and other immunocompromised patients with impaired T cell immunity.

## Results

### Microscale *ex vivo* priming and expansion strategy allows for sensitive gauging of virus-specific T cell immunity

Identification of T cell reactivity in peripheral blood mononuclear cells (PBMCs) by detection of cytokines via ELISPOT or flow cytometry has limited sensitivity due to the low frequency of antigen-specific memory cells in unmanipulated steady-state peripheral blood. This can be partially overcome using a sophisticated approach detecting activation induced markers (AIM) that are upregulated upon stimulation, allowing for sensitive identification of antigen-specific CD4^+^ T helper (Th) cells [30, 31]. However, a large number (0.1-2×10^7^) of PBMCs [24, 32] are required to detect a meaningful signal above the background. Based on our previous experience, we hypothesized that a microscale priming/expansion strategy might unequivocally detect pre-established T cell immune responses against viral antigens even when the frequency of the memory T cells is at the background level[33]. Using overlapping peptide libraries (pepmixes) composed of 15-mer peptides covering full-length immunodominant viral antigens of interest, we directly compared reactivity within PBMC samples (n=8) upon baseline short stimulation (Day 0) and following a 14-day *in vitro* expansion. Baseline PBMCs were evaluated using the AIM method (Fig. S1A upper panel) detecting CD134 and CD137 upregulation [30] and the intracellular detection of TNF-α, IFN-γ and IL-2 (Fig. S1A middle panel). *In vitro* expanded samples from the same donors were tested for the intracellular cytokine secretion upon antigenic re-challenge at day 14 from the start of culture (Fig. 1A, bottom panel). The set of pepmixes included common viral antigens EBV BZLF and EBNA1, Adenovirus (AdV) penton (Ad5), BK virus large T (LT) and VP1 antigens as well as the S1, S2, M and NP antigens from SARS-CoV2, HKU1 and 229E hCoVs. In some donors the antigen-specific cytokine production was detectable upon direct stimulation (Fig. S1B & S1C), while the AIM method revealed reactivity in a larger portion of tested subjects (Fig. S1B & S1C). However, significantly more reactive donors were identified upon 14-day *in vitro* priming/expansion, indicating improved sensitivity of the proposed strategy (Fig. S1B & S1C). Importantly, this approach unmasked reactivity against viral antigens from both tested hCoVs. Whereas direct detection of reactivity on Day 0 (Baseline) by cytokine or AIM (to a lesser degree) was affected by low dynamic range and background signal, the magnitude of the responses in the expanded T cell populations was clearly above background (Fig. S1A), enabling unequivocal identification of reactive T cells. Overall, the micro-scale priming/expansion strategy represents a valid tool for gauging the functional immunocompetence and immune reactivity against the viral antigens of interest, especially when a low frequency of precursors is present and a limited quantity of starting PBMCs is available for analysis.

**Fig. 1:**
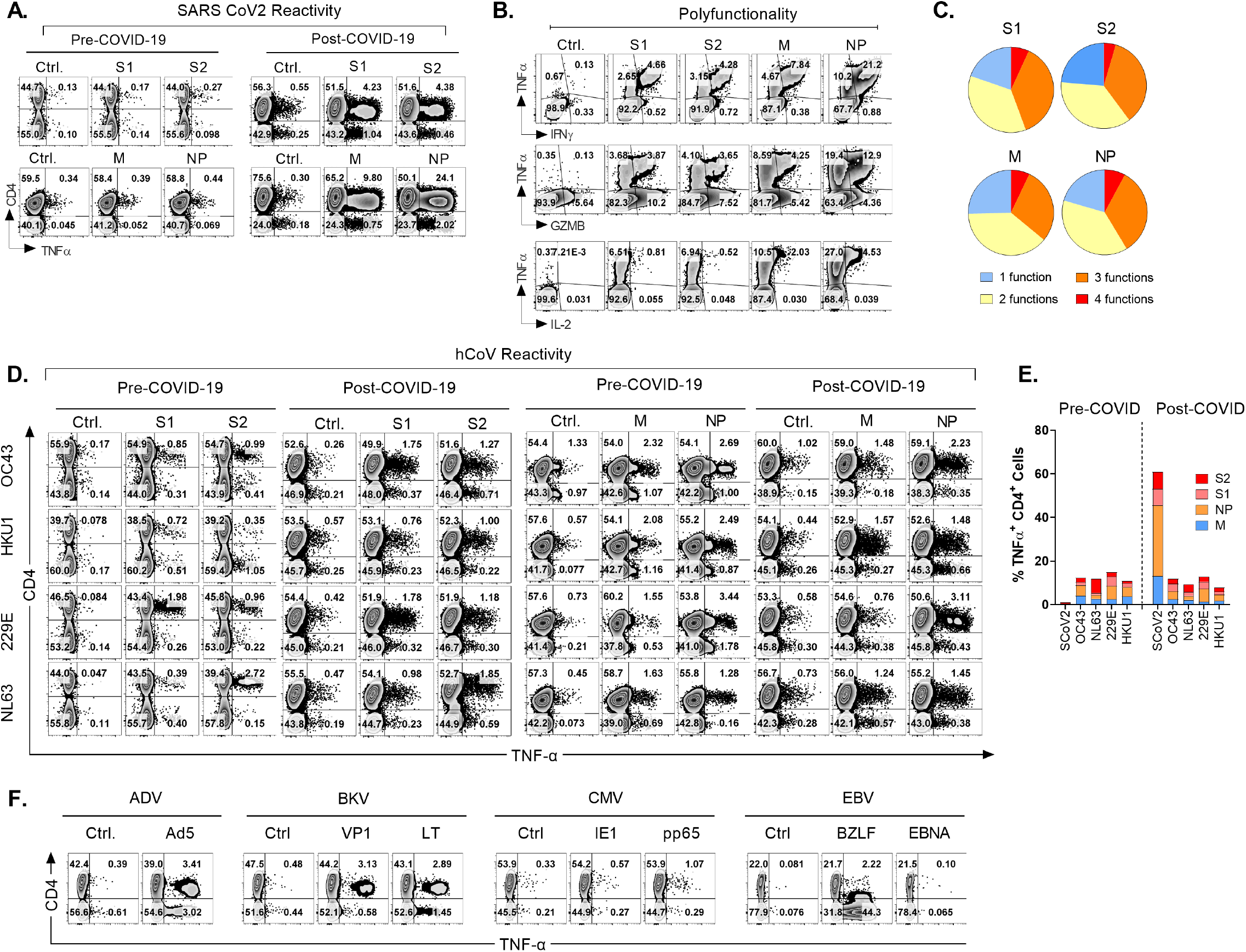
Emergence of potent polyfunctional T cell response following resolution of COVID-19 infection. **A**. Flow cytometric analysis of T cell populations generated upon ex vivo expansion of PBMCs from a healthcare worker (HCW) # 1008 using overlapping peptide libraries derived from indicated SARS-CoV2. PBMCs were collected before and after documented COVID-19 infection. Dot plots were gated on CD3^+^ viable cells and show intracellular production of TNF-α against indicated SARS-CoV2 antigens at the end of 14-day *in vitro* expansion. **B** and **C**. Polyfunctionality analysis of SARS-CoV2 reactive CD4^+^ T cells expanded in vitro from post COVID-19 samples of the same donor. Dot plots were gated on CD3+ CD4^+^ viable cells and show intracellular production of TNF-α vs. IFN-ϒ, vs. GZMB and IL-2 for each indicated SARS-CoV2 antigen. Pie charts show the number of functions (single or multiple types of cytokine production) detected among antigen-reactive cells for each indicated viral antigen. **D**. T cell reactivity in the HCW PBMC samples collected before and after COVID-19 infection expanded upon priming with S1, S2, M, or NP peptide mixes derived from hCoV (OC43, HKU1, 229E or NL63). **E**. Frequencies of SARS-CoV2 specific TNFα^+^ cells among CD4^+^ T cells against indicated peptide mixes as compared to pre- and post-infection reactivity against indicated hCoVs. **F**. Reactivity of ex vivo expanded PBMCs from the same HCW collected pre-COVID-19 primed with peptide mixes derived from the immunodominant antigens of Adenovirus (Ad5), BK polyomavirus (VP1 and LT), CMV (IE1 and pp65) and EBV (BZLF1 and EBNA1) gated identically as in **A**.

### Induction of a broad and robust antigen-specific T cell reactivity against SARS CoV2 in a healthy donor following the resolution of COVID-19

To investigate CoV-specific cellular immunity, we studied serial PBMC samples from healthy volunteers and immunocompromised subjects, with or without known COVID-19 exposure, to analyze T cell immune responses against SARS-CoV2 and related 229E, OC43, NL63 and HKU1 hCoVs. One of the subjects, #1008, a healthcare worker involved in direct patient care, developed PCR-documented COVID-19 approximately three months after initial sample collection, thus allowing for investigation of T cell responses pre- and post-SARS-CoV2 infection. Baseline T cell responses against SARS CoV2 S1, S2, M and NP antigens were minimally detectable above background (Fig. 1A), confirming naïve/unexposed status. In post-COVID-19 samples, a marked increase in reactivity against all four tested SARS CoV2 antigens was observed predominantly within the CD4^+^ T cell compartment. The maximal reactivity was seen against NP antigen with 25.52% TNFα^hi^ cell among CD4^+^ T cells, followed by M (10.29%), S2 (6.14%) and S1 (5.86%; Fig. 1A).

Furthermore, the reactive CD4^+^ T cells displayed a significant polyfunctionality with antigen-specific secretion of TNF-α, IFN-Ỵ, granzyme B (GZMB) and IL-2 with 6.85% of S1 reactive cells, 4.55% of S2 reactive cells, 7.07% of M reactive cells and 7.86% NP reactive cells displayed all 4 functions (Fig. 1B-C). We then tested reactivity to the corresponding immunodominant antigens from the related common α and β-hCoVs in the same donor. The magnitude and pattern of reactivity against all non-SARS hCoV targets pre- and post-COVID-19 remained low and was minimally affected by COVID-19 (Fig. 1D-E). Thus, this HCW acquired robust and highly focused antigen-specific T cell memory against SARS CoV2 post-infection. Moreover, while SARS CoV2-related T cell responses involved mainly CD4^+^ Th cells, *ex vivo* reactivity against antigens derived from the common viral pathogens CMV (pp65 and IE-1), EBV (EBNA1 and BZLF1), BK (LT and VP1) and AdV (Ad5) was seen in either CD4^+^ and/or CD8^+^ T cell compartments of the same subjects (Fig. 1F & Fig. S1D). This suggests that the observed CD4 or CD8 reactivity patterns against each viral antigen depend on the pre-established *in vivo* memory inherent to each virus, and is driven to a lesser degree by the fundamental tendency of the assay to detect only CD4^+^ T cell responses.

### T cell responses against SARS CoV2 antigens among healthcare workers and immunocompromised patients

Next, we analyzed the *ex vivo* T cell responses against the immunodominant antigens of SARS CoV2 in healthy volunteers recruited among healthcare workers (HCW; n=32) and in immunocompromised patients (IP; n=13; table 1). Both unexposed (COVID-; n=25) and exposed (COVID+; n=20) individuals were included. In line with the case presented in Fig. 1, robust *ex vivo* CD4^+^ T cell reactivity targeting SARS-CoV2 S1, S2, NP and M antigens was seen in subjects with documented history of COVID-19 (Fig. 2A; left panel), with a more variable pattern of reactivity observed within the CD8^+^ T cell compartment (Fig. S2A-C). Among COVID+ subjects, M and NP antigens induced a relatively higher frequency of reactive T cells than S1 and S2, indicating that the most vigorous memory was formed against non-spike antigens of SARS CoV2 post-infection. (Fig. 2A, left panel) Antigen-specific reactivity (defined as % TNFα^HI^ cells among CD4^+^ T cells > 0.5%) against SARS CoV2 antigens was also detected in some subjects with no documented COVID-19 disease (Fig. 2A, right panel). Among COVID-unexposed donors collected in the early stages of pandemic, the highest frequency of responses was to the S1/S2 antigens, followed by M and finally the NP antigen, which was the reverse of the pattern observed in COVID-19 survivors. Overall, COVID-exposed subjects displayed a diverse broad pattern of reactivity with a relatively dominant contribution of non-Spike NP and M-specific activity (Fig. 2B; left panel). In contrast, among COVID-unexposed subjects with anti-SARS CoV2 reactivity, responses were narrower and skewed towards Spike antigens (Fig 2B, right panel). Overall magnitude of *ex vivo* SARS CoV2 antigen responses, as measured by frequency of TNF-α^HI^ antigen-reactive CD4^+^ T cells, was significantly higher in COVID-exposed subjects than in COVID-unexposed subjects (Fig. 2C). Representative flow cytometry data of antigen-specific T cell responses in COVID+ subject is shown (Fig. 2D; upper row), as compared to COVID-donor with a relatively robust antigen-specific cytokine release (Fig. 2D, middle row) and a COVID-subject with minimal ability mount *ex vivo* T cell response (Fig. 2D, lower row). Overall, response to at least two antigens was seen in 100% of studied COVID-19 survivors, with 90% of subjects displaying reactivity against at least 3 out of 4 SARS-CoV2 antigens, and 85% of subjects robustly responding to all four pepmixes (Fig. 2E). In contrast, 5 out of 25 studied COVID-unexposed unvaccinated donors (20%) did not show *ex vivo* reactivity (defined as 0.5% above background) against any SARS CoV2 antigens, 8 subjects (32%) mounted a clear response to a single viral antigen, six subjects (24%) showed reactivity against 3 antigens and one subject (#1018) displayed robust recognition of all four SARS CoV2 antigens (Fig. 2F, upper row). Subject #1018 was a HCW who was quarantined in the early stages of COVID-19 pandemic (March 2020) upon developing self-limiting non-febrile upper respiratory tract illness following close contact with a confirmed COVID-19 patient. This subject repeatedly tested negative by PCR and serological studies. Similarly, another subject with multiple myeloma demonstrated a robust T cell response against three SCoV2 antigens (Fig. 2F, lower panel). These, data suggest that both subjects might have developed a mild infection without seroconversion, but with emergence of vigorous T cell responses.

**Fig. 2:**
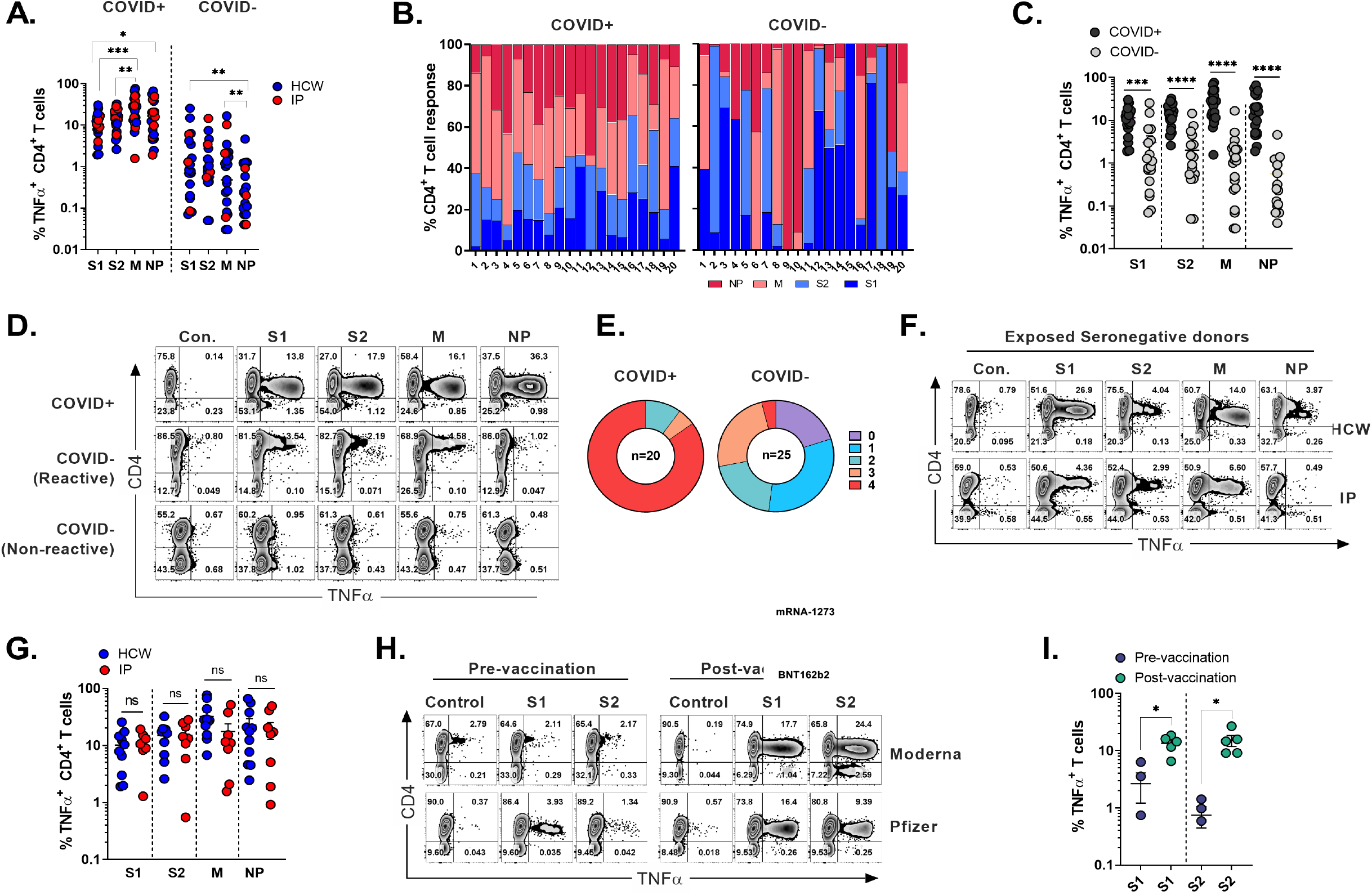
SARS-CoV2 specific T cell responses among HCW and immunocompromised patients with and without history of COVID-19. PBMC samples were collected from healthy volunteers and immunocompromised (IP) subjects exposed and unexposed to SARS-CoV2 and expanded in vitro for 14 days upon priming with SARS-CoV2 peptide mixes S1, S2, M and NP. Reactivity (%TNFα+ T cells) was evaluated in the final cultures upon re-stimulation with cognate peptide mixes. **A**. Frequencies of TNFα-secreting cells among CD3^+^CD4^+^ T cells recognizing indicated SARS-CoV2 peptide mixes in ex vivo expanded cultures. Magnitude of reactivity against peptide mixes in COVID+ and COVID-cohorts was compared using Wilcoxon matched-pairs signed rank test. **B**. Relative contribution of all four SARS-CoV2 proteins to total CD4^+^ T cell response against SARS-CoV2 antigens in individual donors is shown in samples from COVID+ (n=20) and COVID- (n=25) donors. (5 COVID-donors did not show response to any of the 4 antigens.) **C**. CD4^+^ T cell reactivity against indicated SARS-CoV2 peptide mixes among COVID+ and COVID-donors. Statistically significant differences of reactivity between two groups were analyzed by Mann-Whitney test. **D**. Representative flow cytometric zebra plots showing patterns of intracellular TNFα secretion in ex vivo expanded samples from COVID+ donor as compared to COVID-donors with relatively high vs no reactivity upon re-stimulation with SARS-CoV2 peptide mixes. Samples were gated on viable CD3^+^ T cells, unrelated peptide mix was used as negative control. **E**. Percentage of COVID+ (n=20) and COVID- (n=25) donors recognizing (frequency above 0.5%) one or more SARS-CoV2 antigens. **F**. Recognition of indicated SARS-CoV2 peptide mixes in cultures from a PCR negative and seronegative HCW (#1018) with highly probable exposure history (upper panel) and a seronegative immunocompromised patient (#1032) mounting robust T cells response against SARS-CoV2 (lower panel). **G**. Comparison of SARS-CoV2 specific CD4^+^ T cell responses (TNFα^+^) between COVID+ HCW and IP by Mann-Whitney test. **H**. Representative zebra plots demonstrating antigen-specific T cell responses against S1 and S2 CoV2 antigen pre- and post-vaccination. **I**. Frequencies of SARS-CoV2 S1 and S2 reactive CD4^+^ T cells in otherwise unexposed HCWs pre- and post-vaccination. Statistically significant differences of reactivity were determined by Wilcoxon matched pairs signed rank test. Horizontal lines indicate the mean ± SEM. *P < 0.05, **P < 0.01, ***P < 0.001.

Overall, there was no statistically significant difference in the magnitude of CD4^+^ T cell responses directed against all tested SARS CoV2 antigens between HCW and IP COVID-19 survivors, albeit with a trend towards more potent reactivity specific for M and NP antigens among healthy subjects (HCWs), but this analysis might be limited by the small sample size of IP COVID+ survivors (Fig. 2G). Within the CD8^+^ compartment a trend towards lower responses in convalescent IP subjects was seen as well, while more vigorous, although highly variable antigen-specific CD8^+^ responses were mounted by COVID+ HCWs, again, not reaching statistical significance (Fig. S2D).

Finally, robust T cell responses against both subunits of spike protein emerged in the donors who received either BNT162b2 or mRNA-1273 vaccines (Fig. 2H). Vaccination induced a significant increase in frequencies of S-specific T cells when compared to pre-vaccination samples from the same donors (Fig. 2I). Thus, natural infection with SARS-CoV2 or vaccination typically induces T-cell immunity against COVID-19.

### T cell recognition of the divergent epitopes from the emerging variants of SARS CoV2 is largely preserved

The emergence of SARS-CoV2 variants such as delta and omicron BA.4 and BA.5 raise concerns of immune escape that has been well documented for humoral immunity[27, 34]. To examine if previously acquired natural or vaccination-induced T cell immunity cross-reacts against SARS-CoV2 variants, PBMCs from COVID+ (n=6) or unexposed (n=6) subjects who received one of three approved SARS-CoV2 vaccines were expanded *in vitro* using full length ancestral S1/S2 pepmixes. Resulting T cells were then tested for reactivity against peptide pools of 8 SARS-CoV2 variants of concern (VOCs) spanning mutated regions of S protein as compared to the matching S antigen pools from the ancestral (wild type; WT) virus. Response to each variant pool was expressed as stimulation index (SI), defined as fold change (FC) in frequency relative to negative control and then compared to WT-counterpart.

Among COVID-exposed donors the overall SARS-CoV2-specific CD4^+^ T cell response against all 8 variants was largely preserved (S.I range 1.14 to 0.53 (+/-SEM); Fig. 3A). Mean reductions of 27.87%, 22.5% and 16.17% were observed against *beta, epsilon* and *gamma* variants respectively, while only a marginal decline was seen against *alpha* (8.5%), *delta* (5.17%) and *kappa* (0.83%) variants. The greatest mean reduction of 47% was observed in case of *omicron*, consistent with the high mutation burden of this variant. Interestingly, a higher mean CD4^+^T cell response (S.I 1.14) was seen against the eta variant.

**Fig. 3:**
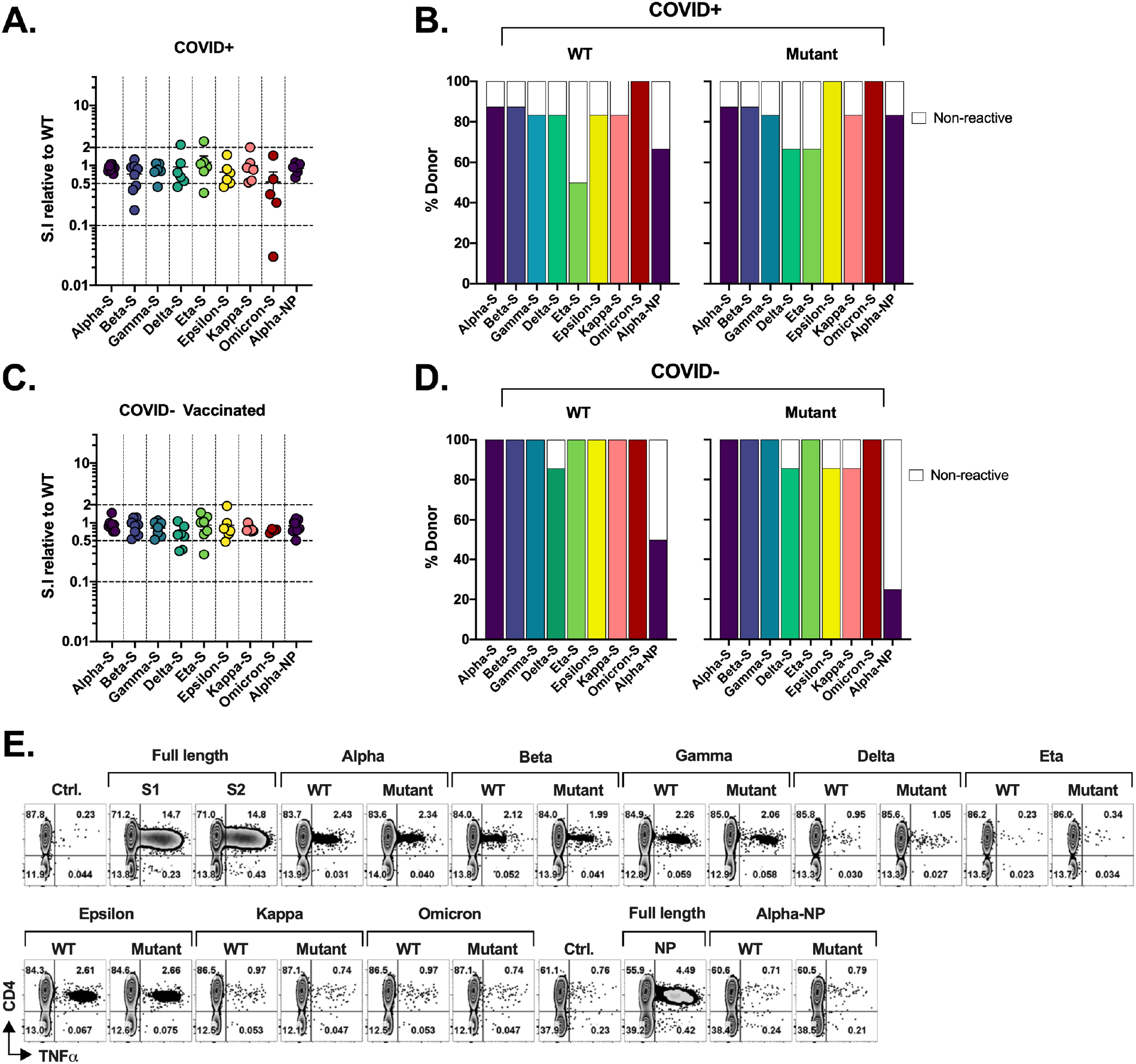
SARS-CoV2 specific response in HCWs and IP patients. T cell cultures primed against SARS-CoV2 Spike (S1/S2) and NP peptide mixes were generated from COVID-19 survivors (COVID+) and unexposed fully vaccinated subjects (COVID-). **A**. Relative response against variant antigen pools as compared to their WT counterpart in COVID+ donors expressed as stimulation index (S.I). **B**. Proportion of cultures displaying unequivocal reactivity against WT (left panel) and Mutant (right panel) peptide pools of indicated SARS-CoV2 variants among COVID+ donors. **C**. Relative response against indicated variant antigen pools as compared to their WT counterpart in COVID-donors expressed as S.I. **D**. Proportion of samples from COVID-donors showing reactivity to WT (left panel) and Mutant (right panel) peptide pools of indicated SARS-CoV2 variants. **E**. Representative flow cytometric analysis illustrating antigen-specific intracellular TNFα secretion upon stimulation with SARS-CoV2 variants as compared to the counterpart WT peptide pool in S and NP cultures from a vaccinated HCW with no history of COVID-19.

At an individual level, maximum of 33-fold reduction was observed in a single COVID+ IP subject against omicron variant, while two additional donors showed more than 2-fold reduction (Fig. 3A). No donors showed more than 10-fold reduction in magnitude of CD4^+^T cell response towards other variants. 3 donors showed more than 2-fold reduction (but less than 10-fold; S.I range 0.18-0.46) in reactivity against *beta*. Relatively robust cross-reactivity was seen against all the other variants tested. Notably, some COVID+ donors demonstrated more than 2-fold increase in response to the variants compared to ancestral pool, namely one donor each against *delta, eta*, and *kappa* variant, possibly due to infections with these variants. Overall, all tested COVID+ survivors that showed reactivity towards the WT-peptide pool also cross-recognized all other variants except *delta* which showed fewer reactive donors compared to its WT counterpart (5/6 vs 4/6 or 83.33% in WT vs 66.67% in delta variant) but the observed magnitude of cross reactivity was typically lower against variant (S.I <1.0) than the WT counterpart (Fig. 3A-B). Curiously, fewer subjects recognized WT pools than the mutants of *eta* or *epsilon* variants (4/6 vs 5/6 or 66.67% vs 83.33% and 5/6 vs 6/6 or 83.33% vs 100% respectively).

In the vaccinated COVID-group (Fig. 3C), mean CD4^+^ T cell S.I was 0.9413 to 0.6414 relative to WT counterpart, with maximum mean reduction of 35.86% seen against *delta* variant followed by 25.75% mean reduction towards *omicron. Kappa, gamma*, and *beta* variants displayed mean reduction of 21.14%, 18.29% and 11.12% respectively. Less than 10% of mean reduction in CD4^+^ T cell response was observed in case of *alpha* (5.87%), *eta* (8.14%) and *epsilon* (9.14%) variants.

At an individual level, only one donor failed to cross-react with *epsilon* and *kappa* pools, while retaining the reactivity towards the control WT peptide pools (Fig. 3D). None of the other donors displayed more than 10-fold loss of response compared to WT peptide pool apart from the *epsilon* and *omicron* variants, with maximum of 3.44-fold reduction observed in a vaccinated HCW against eta variant. Only 2 other HCW donors showed more than 2-fold reduction against the *delta* variant (3-fold and 2.86-fold reduction).

As expected, few vaccinated COVID-donors displayed any reactivity to WT and equivalent *Alpha*-NP pools, while responses to NP WT and mutant pool were easily detected in COVID-exposed donors with broad post-infectious immunity. Fig. 3E shows a representative T cell response of an unexposed (COVID-) vaccinated HCW subject to all the variants tested. Importantly the responses measured here are against the variant epitopes spanning only divergent regions (Fig. S3A) and therefore represent only a minor portion of the overall T cell immunity against the full-length Spike proteins. Notably, among COVID-19 survivors the T cell repertoire targets full length viral antigens much more extensively (see Fig. 2) than the relatively limited responses directed towards the small number of variant-specific epitopes. Similarly, total anti-Spike T cell reactivity induced by vaccination (Fig. 3E) is much more comprehensive. Thus, the broad T cell immunity against SARS CoV2 might provide substantial cross-protection against emerging variants, as it targets full length of the immunodominant viral proteins.

### Cross-reactivity is partially preserved between Spike-specific T cell responses against the ancestral and *Omicron* variant

The recent dominant *Omicron* variant harbors the highest number of amino acid alterations (37) within the Spike protein as compared to previous variants. Furthermore, omicron is more transmissible than previous variants, evades neutralizing antibody responses and has greater capacity for reinfection.

Recent studies have examined T cell responses to omicron BA.1 spike antigen [35, 36]. However, the ability of omicron S-specific T cells to cross-recognize S protein of ancestral virus or other variants has not been studied in detail. Thus, we performed a comprehensive comparison of T cell responses to the omicron variant and the ancestral virus. We used full-length S1/S2 peptide libraries of omicron variant and ancestral viruses to prime PBMCs of COVID+ or vaccinated COVID-unexposed subjects. Resulting T cell populations were tested for recognition of cognate and counterpart full-length S1/S2 peptide mixes as well as for the ability to cross-recognize VOCs.

In all tested donors (n=10), there was no statistically significant difference in frequency of S-reactive (TNFα^+^) T cells upon *ex vivo* expansion with omicron and the ancestral S1/S2 antigens (Fig. 4A-B); albeit the reactivity was significantly reduced upon cross-stimulation as compared to the cognate antigen stimulation (Fig. 4C). The overall magnitude of the ancestral S-specific T cell response when challenged with *omicron* showed a mean loss of 52.29%, suggesting around 47% of the reactivity is remaining against *omicron* variant. Conversely, overall magnitude of the *omicron* spike-specific T cell responses when challenged with ancestral spike showed a mean reduction of 54.09% compared to response against cognate antigen with approximately 45% of the reactivity retained against ancestral variant.

**Fig. 4:**
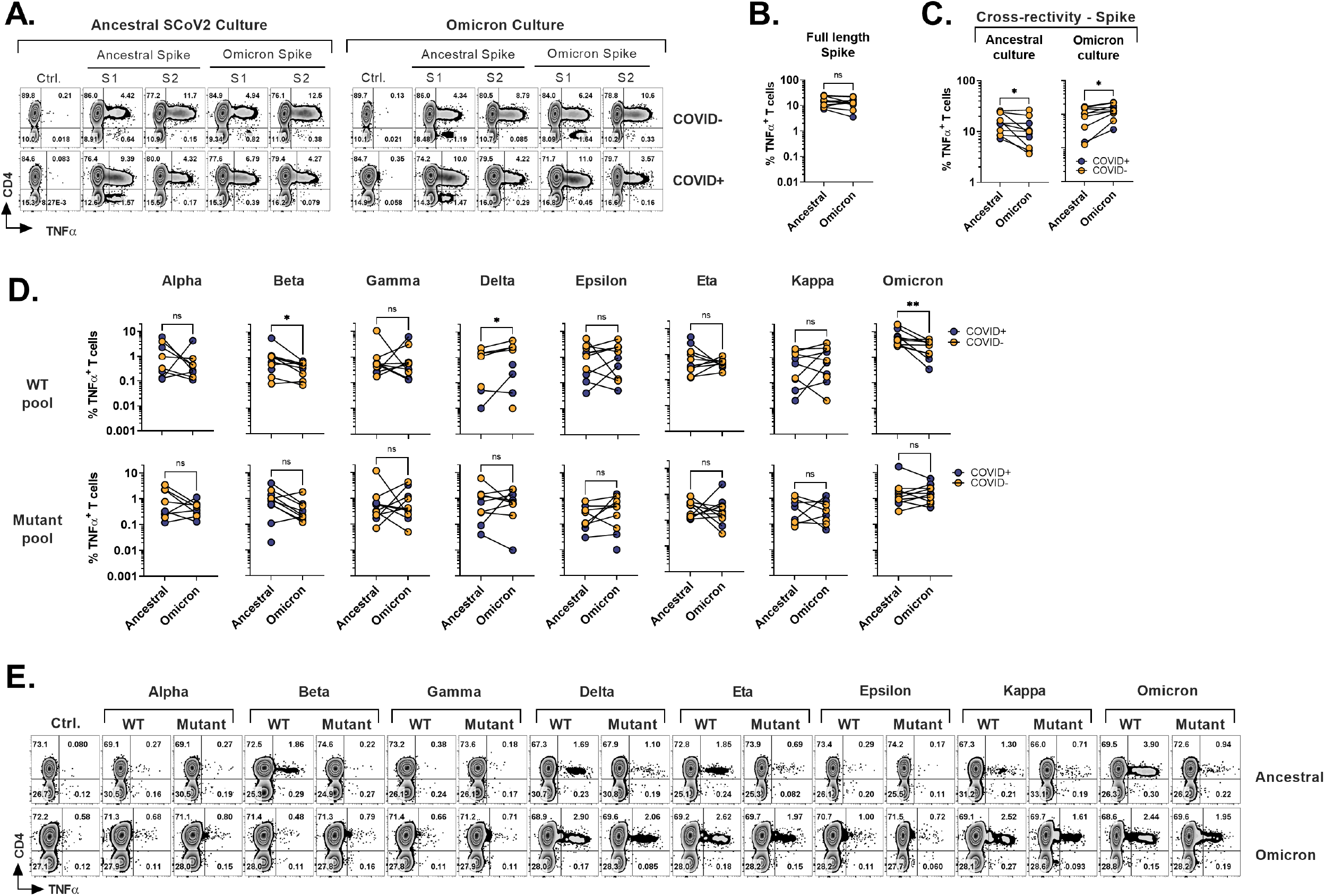
Cross-recognition of T cells specific to ancestral and omicron spike antigens. PBMCs from exposed (COVID+; n=5) and vaccinated but otherwise unexposed donors (COVID-; n=5) were primed and expanded using full-length ancestral and omicron Spike peptide mixes. Reactivity was tested against the indicated antigens. **A**. Representative flow cytometric analysis of antigen-specific reactivity (TNFα secretion) in cultures from COVID+ (#1030) and COVID-vaccinated (#1007) donors depicting T cell response to ancestral and omicron full length spike protein and respective RBD peptide pool. **B**. Frequency of antigen-specific T cells generated against ancestral and omicron S peptide mixes and tested against cognate antigens. **C**. Frequency of cross-reactive T cells between ancestral and omicron Spike cultures, as compared to cognate peptide mix reactivity. **D**. Comparison of cross-recognition of indicated SCoV2 variants in cultures initially generated by priming with full length ancestral or omicron spike peptide mixes. **E**. Representative dot plots showing flow cytometric analysis of cross-recognition of indicated SCoV2 variant epitopes in comparison to the set of matching ancestral (WT) epitopes in cultures initially generated in E. Statistically significant differences of reactivity was determined by Wilcoxon matched-pairs signed rank test. *P < 0.05, **P < 0.01, ***P < 0.001.

To examine whether *omicron*-primed T cells retain the ability to cross-recognize other known variants compared to ancestral spike, we challenged *omicron* or ancestral spike specific T cells with VOCs and VOIs-derived antigens (Fig. 4D-E). All the variant pools used in the study displayed comparable cross-recognition between *omicron* and ancestral S-specific T cells (Fig. 4D, lower panel). Interestingly, the responses against the WT counterpart pools of *beta, delta* and *omicron* variants were lower in *omicron*-primed T cells compared to that of ancestral S-primed cultures (Fig 4D upper row), while these differences were not observed against the mutated set of antigens (Fig 4D lower row). A representative dot plot of a vaccinated donor is shown in Figure 4E.

Additionally, subject #1008 (see Fig. 1) who developed natural immunity during the early stage of pandemic, received full course of vaccination and subsequently developed reinfection during the 4^th^ wave dominated by *omicron* variant (Fig. S3B), allowing for unique longitudinal analysis of CD4^+^ and CD8^+^ T cell responses (Fig. S3C & S3D). Marked S-specific T cell reactivity was consistently seen following the initial infection, the course of vaccination, and re-infection with likely *omicron* variant, but the magnitude of anti-NP and M-T cells responses relatively declined as compared to the primary infection (Fig. S3C & S3D). We then evaluated anti-Spike reactivity of T cell samples from early and late time points upon priming with full length S1/S2 pepmixes of corresponding variants (Ancestral for infection #1 and *Omicron* post-infection #2) and testing for their ability to cross-recognize other variants. (Fig. S3E). The post-reinfection sample displayed enhanced CD4^+^ and CD8^+^ T cell responses against full-length *omicron* pool in contrast to the initial draw, suggesting variant-specific priming. Simultaneously, the responses of both ancestral and *omicron* S-specific T cells against other variants were compared and expressed as S.I (Fig. S3F). As expected, *omicron* S-specific T cells mounted a more potent response to cognate *omicron* sub-pool, but they also produced a relatively higher response against the *kappa* variant (Relative S.I of 0.1346 to 0.8316 respectively). Other variants displayed similar S.I between both ancestral and *omicron* S-specific T cells, indicating preserved cross-reactivity. In summary, the magnitude of response against the *omicron* S1/S2 antigens is reduced by at least half following the COVID-19 or vaccination with ancestral variant. Nonetheless, prior vaccination or previous COVID-19 infection can induce cross-prime against *omicron*.

### T cell responses against common hCoV antigens are seen in COVID-naïve and COVID-exposed individuals

SARS-CoV2 shares the general structure and at least partial sequence homology with the corresponding structural proteins of the endemic hCoVs [24, 37], thus previously acquired immunological memory against those viruses might cross-recognize COVID-19, possibly providing partial protection. Conversely, there might be a broader response to non-SARS hCoVs following the recovery from COVID-19. To better characterize the spectrum of anti-hCoV T cell responses we tested *ex vivo* reactivity using the custom set of pepmixes derived from S1, S2, M and NP antigens of all four endemic hCoVs. A spectrum of antigen-specific activity among COVID-19 survivors and unexposed COVID-subjects was observed, involving both Spike and non-Spike antigens (Fig. 5A-5B), likely representing an imprint of past community-acquired endemic hCoV infections, as illustrated in Fig. 5C depicting recognition of all antigens from multiple hCoVs in a COVID+ donor. Both COVID+ and COVID-donors also displayed vigorous, but highly variable antigen-specific CD8^+^ responses (Fig. S4A-S4B). Overall, reactivity to at least one of the antigens of each hCoVs (Fig. 5D) was found in all tested individuals.

**Fig. 5:**
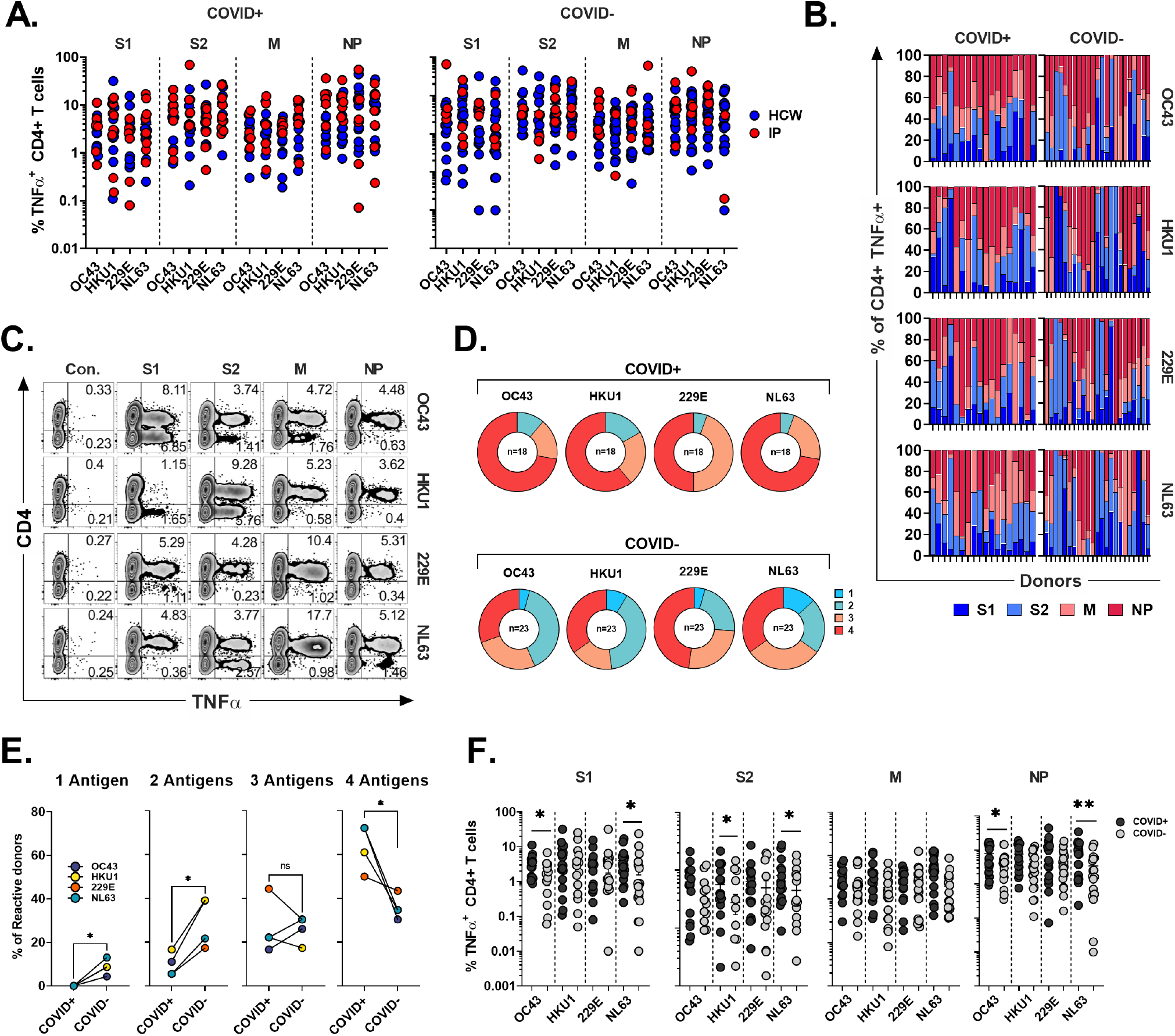
CD4^+^ T cell responses against the immunodominant antigens from “common cold” hCoVs in HCWs and immunocompromised patients. T cell cultures from indicated donors were generated upon priming with peptide mixes for S1, S2, M and NP antigens of α-hCoVs 229E and NL63 and β-hCoVs OC43 and HKU1 and tested as day 14 for reactivity. **A**. Frequencies of antigen-specific CD4^+^ T cells reactive (TNFα+) to S1, S2, M and NP antigens in samples obtained from COVID+ (left panel) and COVID-(right panel) donors. **B**. Relative contribution of reactivity against indicated antigens of each hCoV to total CD4^+^ T cell response in individual COVID+ and COVID-donors. **C**. Representative example of flow cytometric analysis demonstrating broad anti-hCoV response against S1, S2, M and NP peptide mixes in a subject with documented COVID-19 exposure (#1015). **D**. Percentage of donor samples with reactivity against one or more antigens of all analyzed hCoVs among COVID+ and COVID-cohorts. Recognition was defined as frequency of antigen-specific TNFα-secreting T cells >0.5%. **E**. Frequency of donors displaying reactivity against either one, two, three or four antigens (S1, S2, M, NP) of indicated hCoVs in COVID+ and COVID-cohorts. **F**. Statistically significant differences of reactivity were determined by Mann-Whitney test. Horizontal lines indicate the mean ± SEM. *P < 0.05, **P < 0.01, ***P < 0.001.

However, concurrent robust responses (%TNFα^+^ >0.5%) against all four antigens from each hCoV were significantly more prevalent in COVID+ samples than COVID-samples (Fig. 5E). To further examine whether previous SARS-CoV2 infection affects reactivity towards hCoVs, we inspected the magnitude of responses against hCoVs in COVID+ and COVID-donors (Fig. 5F). A significantly higher reactivity was seen against S1 antigen of OC43 and NL63 among COVID-19 survivors as compared to COVID-subjects and against S2 antigens of HKU1 and NL63 with a trend towards increased responses against S2 OC43 and 229E, as well as a significantly higher reactivity against M and NP of NL63 among COVID+ subjects. This was further underscored upon Pearson correlation analysis showing significant correlation between SARS-CoV2 responses and corresponding responses directed against S2 (r^2^= 0.5677) and M (r^2^= 0.6757) of OC43; NP antigens of HKU1 (r^2^= 0.5057) and OC43 (r^2^= 0.6406) (Fig. S4C). When responses against α-hCoVs were analyzed, significant correlation was seen between reactivity directed against S2 of SARS CoV2 and S2 of NL63 (r^2^= 0.5842) and M antigen of 229E (r^2^= 0.4357) (Fig. S4D). Overall, these data suggest a potential link between T cell immunity emerging post-COVID-19 and reactivity potentially directed against other members of CoV family, suggesting possible cross-reactivity and perhaps cross-protection. Indeed, certain COVID-19 survivors mounted broad and robust anti-CoV T cell responses directed against majority of antigens from all analyzed hCoVs.

### T cell responses against hCoV antigens display broad cross-reactivity, but responses against SARS CoV2 in COVID-19 survivors are distinct and specific

To directly evaluate the degree of T cell cross-reactivity against immunodominant antigens from SARS-CoV2 and hCoVs, we primed and expanded PBMCs from donors (n=6, four HCW and two IP subjects) with documented history of COVID-19 using the panel of Spike (S1 plus S2), NP and M pepmixes derived from each hCoV pathogen. The resulting T cell populations from each donor were tested against the cognate pepmix initially used for priming and for cross-reactivity with the counterpart antigens from the other members of CoV family, generating a matrix map of potential cross-reactivity (Fig. S5A). Two representative examples (#1015 and #1023) are shown in Figure 6A, while all analyzed donors are summarized in Figure 6B as a heatmap distribution indicating antigen-specific CD4^+^ T cell reactivity against all 5 viruses (Fig. 6B), where the horizontal axis represents the virus and antigen used for priming and expansion of the T cell cultures while vertical axis represents the virus and the antigen with which the T cell cultures were stimulated. Subject #1015 was a patient with active multiple myeloma who developed symptomatic COVID-19 at the beginning of the pandemic (March 2020) following a cycle of chemotherapy and subsequently underwent autologous SCT. Subject #1023 is a patient with history of erythropoietic protoporphyria (EPP), failed liver and hematopoietic stem cell transplant followed by the second liver and SCT rescue who experienced COVID-19 and was previously described in a case report[38].

**Fig. 6:**
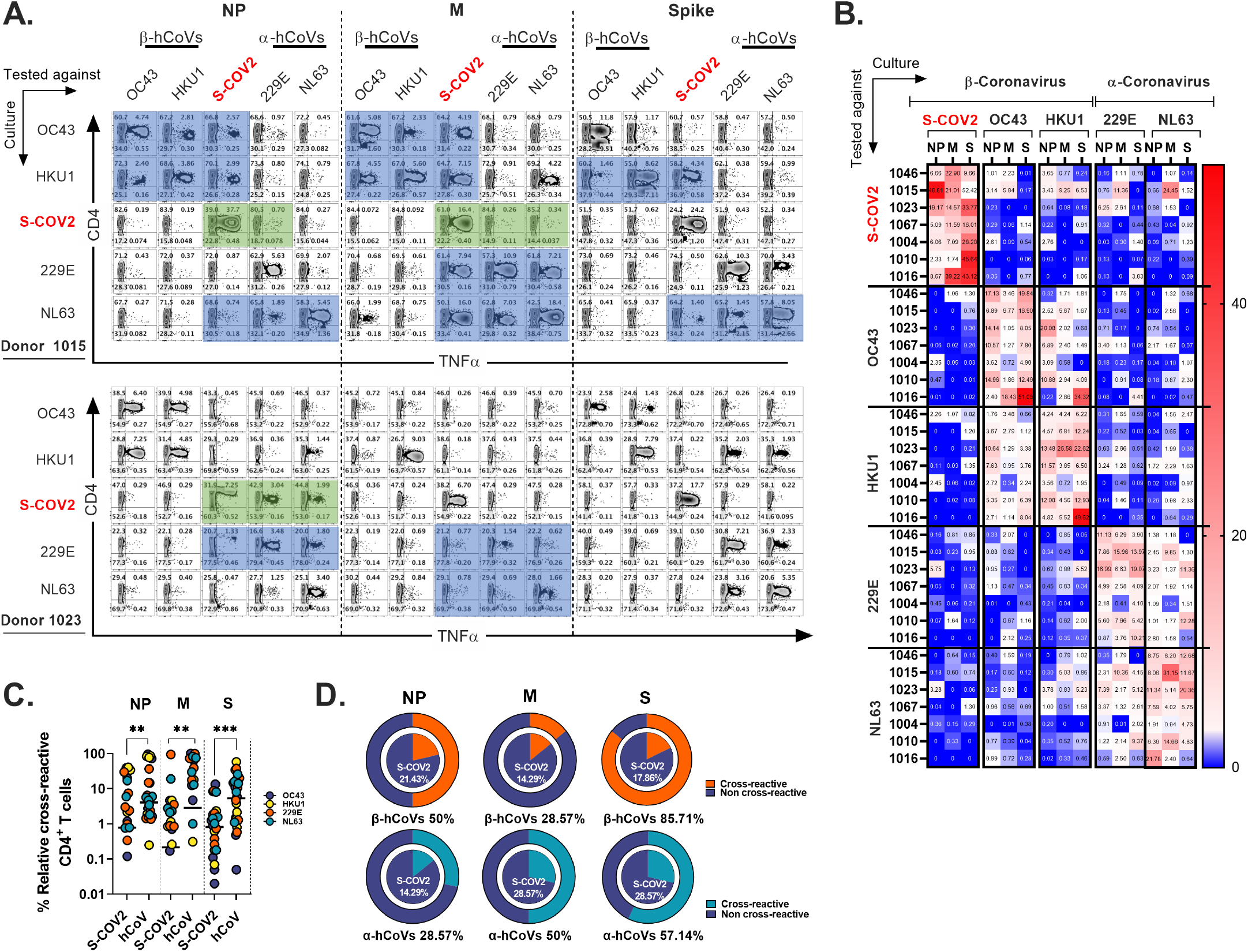
T cell cross-reactivity between SARS-CoV2 and hCoVs. Comprehensive T cell cross-reactivity analysis among two α-hCoVs, two β-hCoVs and ScoV2 against three immunodominant antigens S, M and NP. PBMCs from each donor were stimulated by either S1 & S2 or M & NP peptide libraries against all 5 viruses generating 10 culture conditions per donor. Each culture was challenged by its counterpart antigen from other four viruses along with cognate antigen creating a cross reactivity matrix. **A**. Representative dot plots showing full cross-reactivity panel between five coronaviruses against three antigens. **B**. Heatmap showing relative cross-reactivity in CD4^+^ T cell compartment. **C**. Comparison of relative cross-reactivity between SCoV2 and hCoVs against M, NP, and S protein among CD4^+^ T cell compartment. Horizontal lines indicate the mean value. **D**. Relative frequency (%) of donors displaying cross-reactivity between ex vivo expanded T cell populations specific for SARS-CoV2 (S-CoV2; inner circle) and indicated α-and β-hCoVs (outer circles) against M, NP, and S protein within the CD4^+^ T cell compartment. Statistically significant differences of reactivity were determined by Mann-Whitney test. **P < 0.01, ***P < 0.001.

The *ex vivo* expanded T cells from both donors vigorously recognized cognate antigens of SARS CoV2 and the majority of hCoVs. Cross-recognition was clearly present between counterpart antigens from the closely related non-SARS α- and β-hCoV (NL63 vs 229E and OC43 vs. HKU1, respectively), as anticipated by the relatively high degree of homology between these viruses (Fig. 6A). In subject #1015, β-hCoV (OC43 and HKU1) NP- and M-specific T cells recognized NP and M pepmixes of SARS-CoV2, indicating cross-reactivity (Fig. 6A). Furthermore, hCoV HKU1 S-specific T cells cross-recognized SARS-CoV2 S antigen. This pattern was also seen in α-hCoV specific T cell responses directed against M antigen, and to a lesser degree against NL63 S and NP antigens where some cross-reactivity with SARS-CoV2 is visible. However, while this donor mounted highly specific and potent response against NP, M and S antigens of SARS-CoV2 (49.15%, 21.19% and 50% CD4^+^ TNFα^HI^ T cells respectively), these T cells displayed minimal ability to cross-recognize related antigens from other hCoVs (Fig. 6A, upper panel with green boxes). In subject #1023, upon priming with NP SARS-CoV2 18.52% of CD4^+^ T cells recognized cognate NP and cross-recognized counterpart NP of α-hCoV 229E and NL63 (6.6% and 4.25% TNFα^HI^ T cells respectively; Fig. 6A lower panel with green box). Reciprocally, there was a cross-recognition between 229E NP specific cells and SARS CoV2 NP specific cells (17.33% cognate vs 6.6% for SARSCoV2 CD4^+^ TNFα^HI^ T cells). However, SARS-CoV2 M and S-specific T cells displayed little cross-reactivity against the counterpart pepmixes from the other hCoVs (Fig. 6A, lower panel blue boxes). Extensive cross-reactivity between two related α-hCoVs (229E vs NL63) and two β-hCoVs (OC43 and HKU1) was seen in samples from all donors, while SARS-CoV2 responses displayed relatively little cross-recognition (Fig. 6B). These observations suggest that upon resolution of COVID-19 the emerging T cell responses against SARS CoV2 NP, M and S antigens are highly focused and distinct from the responses mounted against the related non-SARS α and β hCoVs. However, upon priming with non-SARS hCoV antigens a degree of non-reciprocal cross-reactivity against SARS-CoV2 antigens may be observed in the same subjects, that is likely targeting different non-dominant epitopes (Fig. 6A-B). Consequently, T cells expanded using antigens from non-SARS hCoVs showed significantly higher relative cross-reactivity toward the counterpart SARS CoV2 targets while SARS-CoV2-specific T cell cultures were relatively less cross-reactive with non-SARS CoV2 antigens in the same donor (Fig. 6C). Additionally, more donors displayed cross-reactivity (Fig. 6D) in T cell cultures primed initially with non-SARS β-hCoV and α-hCoVs antigens (external pie charts) as compared to only few samples displaying cross reactivity in the opposite direction (internal pie charts). Overall, the data indicate relatively high degree of cross-reactivity between T cells specific for related non-SARS hCoV family members, while variable and non-reciprocal cross-reactivity might be seen with the antigens of SARS-CoV2. These observations might be of practical use when designing strategies aimed at development of universal anti-CoV vaccination or cell-based immunotherapy.

### Distinct T cell receptor (TCR) repertoire targets SARS-CoV2 and common hCoV antigens in survivors of COVID-19

Based on the observed cross-reactivity pattern in flow cytometry, we hypothesized that the focused T cell repertoire directed against SARS CoV2 antigens emerges following the resolution of the infection, with relatively little overlap with the repertoire directed against the antigens from non-SARS α- and β-hCoVs. Conversely, T cells primed to recognize hCoV antigens might be capable of targeting some epitopes from SARS-CoV2, but this cross-recognition favors distinct antigens and is dominated by a non-overlapping clonotypes. To further characterize this phenomenon, in-depth analysis of the TCRβ repertoire of T cells specific for SARS CoV2 NP, M and S antigens and their hCoV counterparts was performed in two survivors of COVID-19: IP subject #1023 (previously described) and an otherwise healthy HCW #1046. The strategy used for TCR sequencing of antigen-specific T cells is outlined as a schematic diagram in supplementary figure S5B.

Virus-specific T cell cultures were generated from both donors using the M, NP and S pepmixes from all 5 viruses (OC43, HKU1, SCoV2, 229E and NL63). Antigen-specific T cells were FACS-sorted using TNF-α cytokine capture (Fig. S5C)[39], allowing for analysis of highly purified antigen-specific T cells with known antigenic reactivity and largely devoid of passenger T cells. The CDR3 TCRβ repertoires of the isolated M, NP and S-specific T cells targeting each virus were analyzed using Adaptive Biotechnologies’ ImmunoSEQ platform.

The number of unique TCRβ sequences, defined by CDR3+V+ J at the amino acid level, identified from each subset ranged from 462 to 2849 for donor #1023, and 214 to 1438 for donor #1046, respectively (Fig. S5D). The TCRβ repertoire specific for each antigen displayed high clonality scores (0.14 to 0.68) and low R20 score (range 0.00089246 to 0.015151), indicating dominant contribution of clonal expansion with restricted diversity (Fig. 7A). Top 10 most prevalent SARS-CoV2-specific T cell clones against different antigen epitopes from subject #1023 represented 11.89 to 21.44% of all sequences identified. In Donor #1046 this fraction was 89.53, 82.03, 61.01% of NP, M and S-specific sequences respectively (Fig. S6A) with the most abundant NP-derived sequence representing approximately 60% of the repertoire, indicating a highly reactive clonotype.

**Fig. 7:**
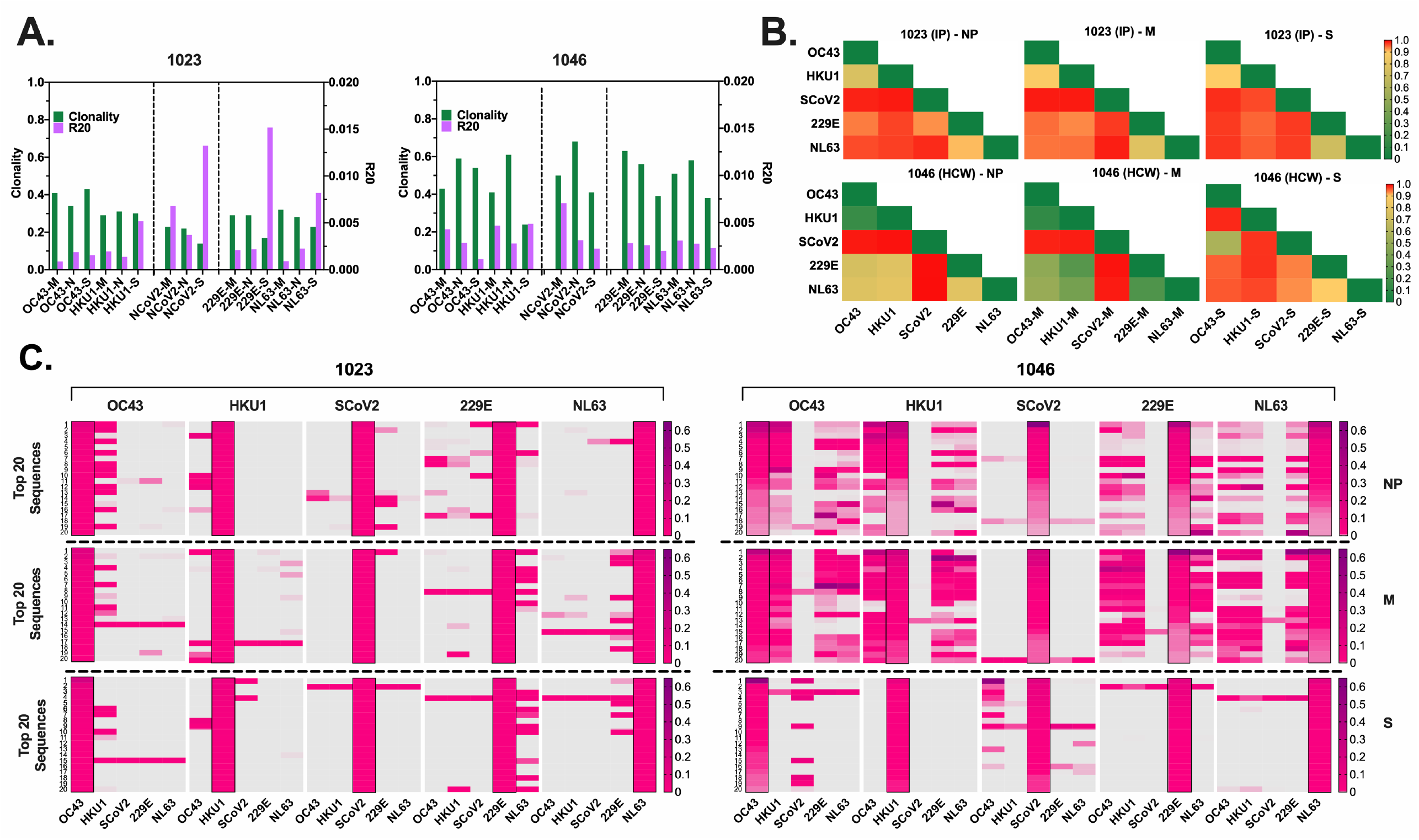
TCR repertoire analysis shows focused response to SARS-CoV2 with few cross-reactive TCRs. **A**. Clonality and R20 values within each antigen-specific T cell populations as described in A. **B**. TCR repertoire similarity between antigen-specific populations recognizing NP, M and S targets from all 5 tested CoVs measured by JSD index (ranges 0-1) and shown in heatmaps. **C**. Heatmaps showing the top 20 dominant clones ranked by frequencies within each designated sample highlighted in black rectangular within each panel, which are specific for NP, M and S antigens from each virus and their cross-reactivity with other CoVs reflected by the co-presence of the same sequence in the same row in donor #1023 (Left panel), Donor 1046 (Right panel).

To estimate the possible presence of the cross-reactive TCRs, we performed global similarity analysis of TCRβ repertoire specific for each antigen of each CoV using Jensen-Shannon divergence (JSD) index. Lower JSD scores (indicating higher repertoire convergence) were seen between the corresponding NP- M- and S-reactive T cells specific for closely related 229E/NL63 α-hCoVs and OC43/HKU1 β-hCoVs (Fig. 7B). There was also some degree of repertoire similarities between populations targeting the corresponding antigens of α- and β-hCoVs, especially in #1046 NP and M samples (JSD ranges from 0.267747 to 0.80066). In the same subject a high similarity was observed between OC43-S and SARS CoV2-S specific T cell repertoire (JSD=0.585714). Lower similarity was seen in both tested donors when SARS CoV2-specific repertoire was compared with the repertoires specific for the corresponding antigens from α- or β-hCoVs, as predicted upon *in vitro* antigen cross-reactivity pattern.

Next the clonal composition of the repertoire was analyzed. In both donors, the majority of isolated TCR sequences were unique for each antigen of each CoVs, as shown in the global Venn diagrams (Fig. S6B), but a small number of TCR sequences shared among all five tested repertoires was seen (ranges 8-30). The analysis of the top 20 most abundant CDR3 sequences specific for each viral antigen (Fig. 7C) revealed certain dominant clonotypes shared between both α- and both β-hCoVs. We also found abundant TCR clonotypes specific for all four non-SARS hCoVs, especially among NP and M-reactive population in subject #1046. However, in both donors there were relatively few dominant TCR sequences derived from the SARS-CoV2 specific population that were shared with other hCoVs, with the exception of #1023 NP antigen overlapping with some sequences from 229E hCoV (Fig. 7C, left panel). In #1046 the overlap was seen predominantly with S antigens of OC43 hCoV (Fig. 7C, right panel) where the most abundant CDR3 sequence (53.1%) was also the most abundant clonotype within the SARS-CoV2-S, representing 14.1% of total CDR3 repertoire, suggestive of possible cross-reactivity. Furthermore, multiple TCRs specific to SARS CoV2 antigens identified from both donors were found in the Adaptive Biotechnologies’ ImmuneCODE database [ref] (Fig. S6C), implying shared/public anti-viral TCRs that are frequently present in the general population. Interestingly, several of the TCR sequences isolated by us from the non-SARS hCoV cultures were found in the ImmuneCODE database. These TCR sequences may represent public clonotypes specific against “common cold” hCoVs antigens, rather than uniquely induced by the COVID-19-related antigens. Taken together, a highly focused TCR repertoire emerges in COVID-19 survivors that has relatively little overlap with T cells induced upon exposure to M, NP and S antigens from non-SARS hCoV family members. In contrast, there is a relatively high frequency of shared TCRs specific to α or β-hCoVs expressed in T cells responding to the closely related members of each hCoV subfamily, as predicted by the high degree of functional cross-reactivity.

### Generation of the universal multi-CoV-specific T cells for adoptive immunotherapy or prophylaxis of COVID-19 and hCoV infections

Finally, based on the *ex vivo* priming strategy, we hypothesized that it would be feasible to generate SARS-CoV2-specific T (SCVST) cells using a clinical-grade procedure compatible with current Good Manufacturing Practice (cGMP) methodology. First, the dedicated COVID-19 products were generated by culturing PBMCs from COVID-exposed donors stimulated with SARS-CoV2 M, NP and S1/S2 peptide mixes pulsed into irradiated autologous PBMCS as antigen presenting cells (APCs). Cultures were maintained for 14 days in G-Rex gas-permeable containers (Fig. 8A). The resulting SCVST cells displayed robust expansion (not shown) and predominantly contained CD4^+^ T cells (>90%). Upon *in vitro* stimulation the SCVSTs recognized all SARS-CoV2 antigens, as well as the S1 fragment representing Receptor Binding Domain (RBD) of the virus (Fig. 8B).

**Fig. 8:**
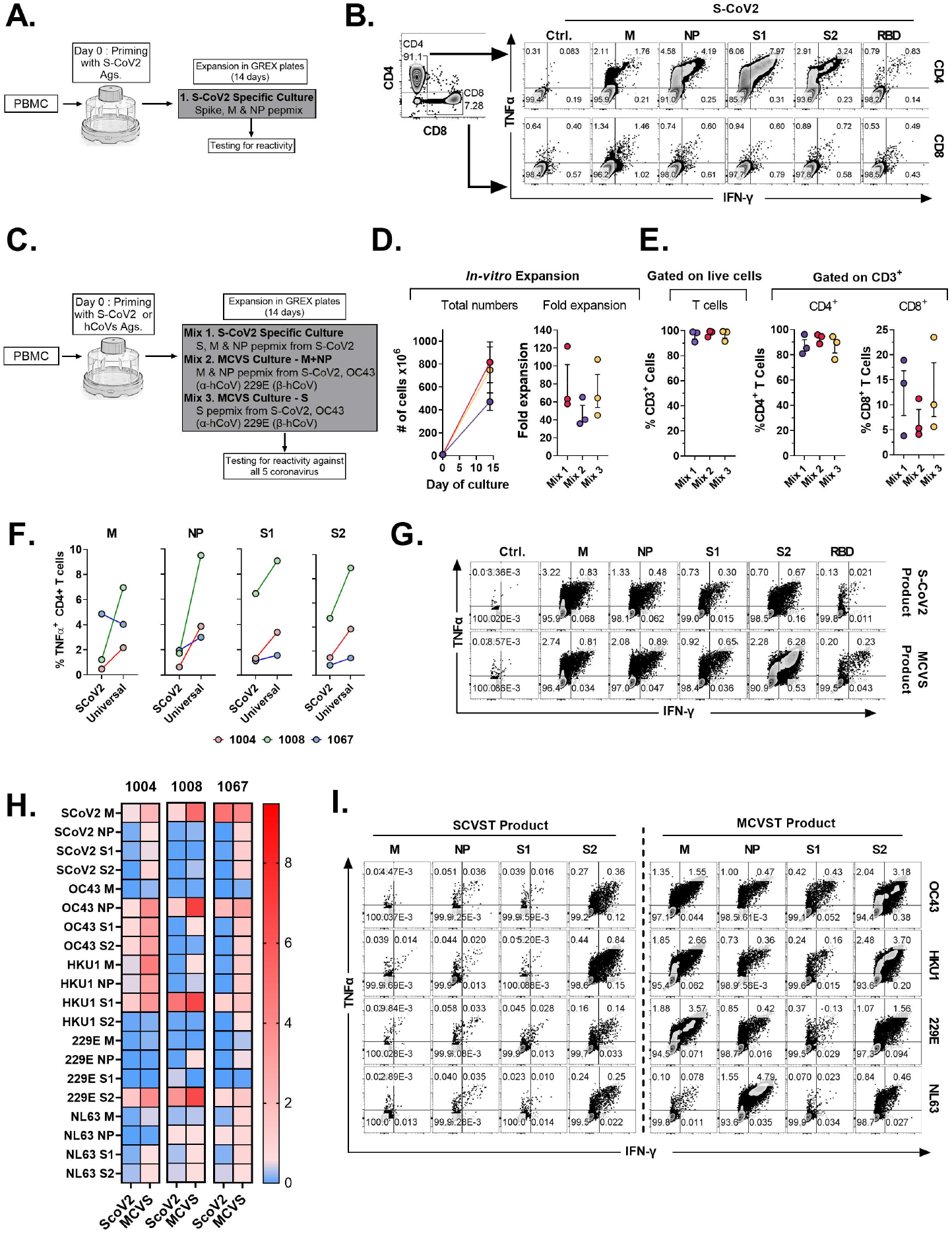
Novel strategy to generate multi-coronavirus specific T cell product for adoptive T cell therapy. **A**. Schematic diagram showing strategy to generate clinical grade SARS-CoV2 specific T cells for adaptive cell therapy using G-REX bioreactors. **B**. Representative dot plots showing reactivity of SARS-CoV2 specific T cell product in CD4 and CD8 T cell compartments. **C**. Schematic diagram showing strategy to generate multi-coronavirus specific (MCVS) T cells. **D**. Ex-vivo expansion (total viable cells) and fold change of the clinical-scale products generated in G-REX flasks in 14 days. **E**. Percentage of CD3^+^, CD4^+^ and CD8^+^ T cells in the final product. **F**. Comparison of reactive CD4+ T cell frequency between SCVST and MCVST products against indicated antigen **G**. Representative dot plots showing reactivity of SARS-CoV2 specific T cell product and MCVST product from the same donor. **H**. Heatmap showing frequency antigen-specific CD4^+^ T cell against indicated antigens (S1, S2, M, NP) of two α-hCoVs (229E, NL63), two β-hCoVs (OC43, HKU1) and SARS-CoV2. **I**. Representative dot plots showing reactivity of SARS-CoV2 specific T cell product and MCVST product against the indicated human hCoV antigens.

Next, based on the cross-reactivity pattern seen in COVID-exposed donors (Fig. 6), we hypothesized that a broadly applicable universal multi-hCoVs-specific T (MCVST) cell product might be generated by using SARS-CoV2 antigens supplemented with counterpart pepmixes from a single α-hCoV and a single β-hCoV. To test this hypothesis, the parallel cultures were initiated using M, N and S1/S2 antigens of SARS-CoV2 (Mix 1: SCVST cells), a culture primed with M and NP antigens from SARS-CoV2, 229E α-hCoV and OC43 β-hCoV (Mix 2: MCVST cells-M+NP) and a culture primed with S1 and S2 antigens form SARS-CoV2, 229E α-hCoV and OC43 β-hCoV (Mix 3: MCVST cells-S) (Fig. 8C). All cultures expanded robustly (range 35.76-121.88 -fold) yielding 3.6×10^8^ to 1028.6×10^8^ viable cells (Fig. 8D) including over 90% CD3^+^ T cells. Of those over 75% were CD4^+^ (range 76.3 to 95.8%) while CD8^+^ T cells comprised 3.8% to 23.5% of CD3^+^ T cells (Fig. 8E).

Intriguingly, as compared to the dedicated SCVST controls, the universal MCVST cultures contained higher frequencies of SARS-CoV2 M, NP and S1/S2 -reactive cells, although not statistically significant (Fig. 8F) and displayed increased ability to recognize RBD (Fig. 8G). Moreover, the final products also displayed marked ability to recognize non-SARS hCoVs, including the NL63 and HKU1 hCoVs not initially used for priming (Fig. 8H-I). The overall reactivity against all three antigens (S, M, NP) of all 5 CoVs among 3 donors tested is shown as a heatmap (Fig. 8H). Thus, priming with a cocktail composed of antigens from SARS-CoV2 supplemented with one α-hCoV and one β-hCoV antigens generates MCVST cells with enhanced ability to target SARS-CoV2, but also induces broad reactivity against “common cold” hCoVs that frequently affect vulnerable immunocompromised patients. This strategy provides a proof-of-concept to create universal MCVST cells that might have the potential to protect or treat current and future variants and emerging diseases caused by human and zoonotic CoVs.

## Discussion

T cell-mediated responses are essential for efficacy of host defense against viral infections. *Ex vivo* expanded virus-specific T (VST) cells have been successfully used to target common refractory viral infections in immunocompromised patients[40, 41]. There is growing evidence that T cell immunity is also critical for protection against the severe complications of SARS CoV2 infection [42-44]. Notably, patients with inborn defects in B cell functionality, frequently experienced only mild course of COVID-19, suggesting that cellular immunity is sufficient for effective host defenses[45, 46]. Furthermore, immunocompromised cancer patients were relatively protected from severe COVID-19 if T cell function was preserved[18]. Importantly, evidence from the SARS-CoV1 outbreak and emerging findings from COVID-19 indicate that a long-lived T cell memory is induced in survivors[19, 21, 47].

Seasonal infections with “common cold” hCoVs are highly prevalent but remain relatively understudied[9, 48]. Intriguingly, emerging data suggest that pre-existing cross-reactive immunological memory induced by prior infections with endemic hCoVs may ameliorate the severity of the subsequent infection with SARS-CoV2[25]. Understanding the T cell response against SARS CoV2 and related hCoVs is critical for the development of adoptive transfer strategies intended for vulnerable cancer and other immunocompromised patients with impaired cellular immunity. Here we tested T cell immune responses against immunodominant antigens (S, M, NP) from SARS-CoV2 and counterpart antigens from α-hCoVs (229E and NL63) and β-hCoVs (OC43 and HKU1) in exposed and unexposed healthy volunteers and immunocompromised survivors of COVID-19, using a custom panel of overlapping peptide libraries including commonly investigated S1 and S2 proteins, as well as less-studied NP and M proteins from each virus. We used the microscale priming/expansion strategy similar to clinical-grade manufacturing of VST cells, incorporating a period of specific expansion and enrichment prior to antigenic testing, enabling unequivocal detection of antigen-specific T cell repertoires. Our approach requires relatively small number of PBMCs, is highly sensitive and allows for functional determination of the immunocompetence upon re-challenge with viral antigens, permitting for unbiased determination of potential cross-reactivity. We have previously used this method to measure the baseline T cell immunity of patients with Progressive Multifocal Leukoencephalopathy (PML) against John Cunningham (JC) Polyomavirus antigens LT and VP1 as a marker to predict the likelihood clinical response to PD-1 blockade[33]. In contrast, the conventional methods (ELISPOT or AIM) have the advantage of estimating the frequency and phenotype of pathogen-specific T cells directly in peripheral blood, but they require large numbers of cells and may be less sensitive when analyzing low-frequency events[30, 49].

Consistent with previous reports[13, 50], we observed T cells responding to SARS-CoV2 antigens in many unvaccinated/unexposed individuals with no documented history of COVID-19, but these responses were highly variable in magnitude and generally favored Spike antigens. Vigorous anti-S1/S2 T cell responses emerged post-vaccination even in immunocompromised subjects, whereas in survivors of COVID-19, broad T cell repertoire targeting Spike and non-spike antigens, such as M and NP, of SARS-CoV2 emerged. Importantly, potent T cell reactivity was seen not only among the otherwise healthy survivors of COVID-19, but also among high-risk patients with a history of hematological malignancy, SCT and SOT, who survived the infection. However, most of the analyzed subjects, including those in the immunocompromised cohort, experienced a relatively mild course of COVID-19 and only one of them required hospitalization. Similarly, our vaccinated IP group included only few very-high risk individuals undergoing active chemotherapy immediately after the transplant.

The anti-SARS-CoV2 responses were predominantly and consistently seen within the CD4^+^ Th cells and retained polyfunctionality as evidenced by production of TNF-α, IFN-γ, GZMB and IL-2. CD8^+^ T cell reactivity was more variable between donors. This might be partially due to the inherent tendency of the longer (15-aminoacid) peptides used for priming to stimulate T cells via class II MHC [51] [52]. However, CD8^+^ T cell reactivity against other viral antigens from BK, AdV and EBV was often detectable in the same assay, suggesting that the pattern of recognition is virus- and donor-specific [14] CD4^+^ T cells might be critical for favorable outcomes of COVID-19[14] and for vaccine responses[53]. The relative magnitude of T cell responses against non-Spike antigens was comparable to or higher than the reactivities against S1 and S2 in survivors of COVID-19. However, the *in vivo* contribution of S, M and NP-specific T cell responses in protection against SARS-CoV2 is unknown. Our data suggest the potential feasibility of targeting non-Spike antigens in the new generation of vaccines[49, 54]. In fact, it has been previously shown that certain class I-restricted NP responses correlate with less viral replication and highly favorable outcomes[55].

Furthermore, our observations indicate that CD4^+^ T cells induced upon priming with the ancestral SARS-CoV2 S1 and S2 epitopes largely retained the ability to recognize mutated antigens representing variants of the virus that emerged during the evolution of the COVID-19 pandemic. This underscores the more promiscuous nature of class II-restricted T cell responses capable of activation by altered peptide ligands. This cross-reactivity was seen in both COVID-19 survivors and in the vaccinated unexposed group. However, in the case of the *omicron* variant harboring over 30 mutations in the Spike protein, *ex vivo* expanded T cells initially primed using the ancestral or *omicron* S1/S2 retained less than 50% cross-reactivity upon testing against their counterparts.

Thus, previous infection and/or vaccination induces broad immunological memory capable of recognizing S1/S2 proteins from novel variants of SARS-CoV2, but at least partial loss of protection is likely against highly divergent variants such as omicron. However, exploiting and/or inducing additional M and NP-directed T cell immunity might provide additional layer of protection against the emerging threats from future variants if anti-Spike cross-reactivity is diminished.

In parallel, we studied reactivity against the endemic hCoV, observing immune responses in the majority of healthy and immunocompromised subjects. Interestingly, COVID-19 survivors displayed a broader reactivity pattern against the endemic viruses, suggesting possible induction of cross-reactive immunity upon resolution of COVID-19 that is increasingly recognized in literature[56, 57]. *Ex vivo* expanded T cells specific for SARS-CoV2 S, M and NP antigens and the equivalent antigens from hCoVs were further tested for cross-reactivity. The most vigorous reciprocal cross-recognition was seen between T cells initially primed to recognize closely related α-hCoV (229E vs NL63) and β-hCoV (OC43 vs. HKU1), while cross-reactivity between T cells specific for α and β antigens was less prominent. Cross-recognition of counterpart antigens from non-SARS hCoVs and SARS-CoV2 was seen in some donors, often displaying unidirectional/non-reciprocal pattern. Namely, T cells initially primed to recognize hCoV antigens were more cross-reactive, whereas priming with SARS-CoV2 antigens induced minimally cross-reactive populations, likely indicating focused immunological memory skewed towards non-shared dominant epitopes[56]. TCR sequencing of highly purified antigen-specific T cells isolated from two COVID-19 survivors elucidates the basis of this observation, revealing dominant clonotypes shared between α or β-hCoV-reactive populations, while SARS-CoV2-specific T cells contained distinct repertoires. However, this phenomenon might warrant more extensive characterization to draw a broader conclusion, as some universal anti-CoV clonotypes were also identified (data not shown). Finally, based on our cross-reactivity observations, we tested feasibility of creating universal Multi-CoV-specific T cells opening the prospect of application in adoptive cellular therapy or prophylaxis in SCT patients, or immunocompromised patients with protracted COVID-19[29, 58]. Concurrent priming of PBMCs with SARS-CoV2 antigens and counterpart antigens from a single α-hCoV and a single β-hCoV induced a population with enhanced capacity to target SARS-CoV2 antigens and with potent activity against all four endemic hCoVs. This synergy further underscores the interplay between the immune responses directed against multiple related members of the CoV family.

In summary, to our knowledge, this is the first study to comprehensively investigate the cross-reactivity of SARS-CoV2 and hCoVs against all three major immunogenic antigens of coronaviruses, namely S, M and NP. Our data supports the hypothesis that a broadly-specific universal anti-CoV T cell-directed vaccines and cellular therapy products are feasible as preventive or rescue immunotherapies.

## Methods

### Patient Blood Sample Collection and Preparation

Healthy volunteers, primarily healthcare workers and associated individuals, with or without COVID-19 exposure, as well as immunocompromised patients were included after informed consent under an IRB approved protocol. Venous blood was collected for serum into a vacutainer containing no anticoagulant. Serum samples were obtained after clotting by centrifuging 3mL of whole blood at 3000rpm for 15 minutes. Serum aliquots were then stored in -80 ºC freezer. PBMCs were isolated using Lymphoprep™ density gradient medium (STEMCELL Technologies Inc., Canada) for the isolation of mononuclear cells, following the product’s protocol. Briefly, blood was diluted 1:1 with sterile dPBS, layered on Lymphoprep™ (ratio 1:1), and centrifuged 30 minutes at 1600rpm, without acceleration/brake. The PBMCs layer was carefully removed, and cells were washed twice with cytokine-free medium (Suppl Table). PBMCs were counted, and immediately used for cell culture and/or flow cytometry analysis, or frozen using cryopreservation medium with 10% DMSO CryoStor® CS10 (STEMCELL Technologies Inc., Canada) and stored in liquid nitrogen.

### Generation of virus–specific T cells

Virus-reactive T cells were generated using commercially available overlapping peptide libraries against immunodominant viral antigens (S, M and NP), purchased from commercial vendors. Lits of all the peptide pools used in the study are listed in supplementary table 2 along with their suppliers and catalog numbers. hCoVs (HKU1, OC43, NL63, 229E), or common viruses (ADV, BKV, CMV, EBV), were obtained from JPT Peptide Technologies or Miltenyi Biotec. SARS-CoV2 PepTivator® Peptide Pools, including the spike protein (PepTivator® SARS-CoV-2 Prot_S1, Prot_S2, Prot_RBD), the nucleocapsid phosphoprotein (PepTivator® SARS-CoV-2 Prot_N), and the membrane glycoprotein (PepTivator® SARS-CoV-2 Prot_M) were obtained from Miltenyi Biotec. The PepTivator® Peptide Pools are constituted by peptides of 15 amino acid length with 11 amino acid overlap. The peptides were grouped into different pools including pool S (equal amounts of Prot_S1, Prot_S2), pool NP, pool M. M and NP peptide libraries for hCoVs 229E, OC43, NL63 and HKU1 were custom synthesized at 70% purity (Peptides and Elephants; Germany). Cryopreserved PBMC were thawed and pulsed with peptide libraries (final concentration of 1μg/ml). After incubation, cells were suspended in CFM (Cytokine Free Media) media with interleukin-7 (10 ng/mL; Peprotech, NJ), and plated on 96 well U-bottom plates. IL-2 (30 IU/mL) was added after 72h. Cells were maintained, fed, and split as needed, every 3 days for approximately 14 days. On day 14, cells were harvested and evaluated for antigen-specificity and functionality by measuring cytokine production and activation marker expression. Cells were restimulated with cognate viral peptides to test their specific reactivity, or with unrelated viral peptides to test their cross-reactivity, via flow cytometry.

For the generation of clinical-grade (GREX®) T cell products, a comparable protocol has been used. PBMCs were pulsed with a master mix of other hCoVs spike S1 and S2 peptide pools, hCoVs membrane and nucleocapsid peptide pools, or SARS-CoV2 spike S1 and S2, membrane and nucleocapsid peptide pools. After incubation, cells were plated in 6 well GREX® (Gas Permeable Rapid Expansion) plates, from Wilson Wolf (Saint Paul, MN), in CFM media with interleukin-7 (IL-7, 10 ng/mL; Peprotech, NJ), and IL-15 (10 ng/mL; Peprotech, NJ). IL-2 (30 IU/mL) was added after 72h. Cells were maintained, fed, and split as needed, every 3 days for approximately 14 days. On day 14, cells were harvested and evaluated for antigen-specificity and functionality. Cells were restimulated with cognate viral peptides to test their specific reactivity, or with unrelated viral peptides to test their cross-reactivity, via flow cytometry.

### Flow cytometry

All antibodies were procured from Biolegend (San Diego, CA) except for Viability Dyes (Miltenyi Biotec, FL). Flow cytometry was performed on PBMCs or cultured cells. Data was acquired on a BD Fortessa, analysis was performed on FACS Diva and FlowJo software (BD Biosciences Corp, San Jose, CA, USA). Antibodies for cell surface markers and intracellular cytokines are listed in Supplementary Table 3. PBMCs were resuspended in 1xPBS with live dead stain (1:200 dilution, Viability Dye, Miltenyi Biotec) for 10 minutes at room temperature. Samples were then resuspended in surface master mix and incubated for 20 minutes at 4ºC. Cells were then washed twice in FACS buffer and fixed for intracellular analysis or resuspended in FACS buffer and acquired at the cytometer.

### Intracellular cytokine staining assay

For intracellular flow cytometry of T-cell clones, cells were stimulated with viral peptide pools to a concentration of 1 mg/mL. 1 mg/mL of brefeldin A (Golgi PLUG, BD Biosciences) and 1 mg/mL of Monensin (Golgi STOP, BD Biosciences), anti-CD28 1ul/mL and anti-CD49d 1ul/mL were added to each well, and plates were incubated for 5h at 37°C 5% CO2. Cells were stained for surface markers following the previously described protocol[38]. Intracellular cytokine staining was performed by following the indication of Fixation/Permeabilization Solution Kit (BD Bioscience), incubating samples with cytokine master mix for 1h at 4ºC. Cells finally resuspended in FACS buffer and analyzed by flow cytometry.

### Activation induced marker assay

Cells were cultured for 24 hours in the presence of indicated antigen pools in 96-wells U bottom plates at 0.5×10^6^ PBMC per well. A stimulation with DMSO was performed as a negative control, and stimulation with a combined Anti-CD3 antibody (OKT3) was included as a positive control. Cells were stained with AIM markers CD137 and CD134[24, 30]. List all the antibodies used in the study is included in Supplementary table 3.

### TCRβ CDR3 DNA sequencing

Virus-specific T cells were first expanded for 14 days as mentioned above. On day 14, the expanded cells were stimulated by cognate antigens in presence of TNFα-PE antibody and TNF-α Processing Inhibitor (TAPI) or 4 hours. Cells were then stained for CD3, CD4 and CD8 surface markers. Antigen-specific T cells defined as TNFα^hi^ were sorted using BD Influx cell sorter at the Columbia Center for Translational Immunology (CCTI) Flow Core. DNA from sorted cells was isolated using Qiagen’s DNA extraction kit (Cat#69504). Purified DNA was measured using nanodrop and sent to Adaptive biotechnologies for TCRβ CDR3 DNA sequencing.

### TCRβ CDR3 sequencing data processing and analysis

DNA was frozen down at -20C and shipped on dry ice to Adaptive Biotechnologies (Seattle, WA) for high throughput TCRβ sequencing. The TCR sequencing data were retrieved from Adaptive’s ImmunoSEQ software. PCR amplification, read sequencing, and mapping, with bias correction and internal controls, were performed by Adaptive Biotechnologies, returning tabulated template counts corresponding to unique bio-identity (CDR3 amino acid sequence + TRBV gene + TRBJ gene) across all samples. Analysis of TCRβ repertoire bulk DNA-seq data was performed in R, Rstudio and Microsoft Excel. Clonality, which ranges from 0 to 1, is primarily used as a measure of diversity, such that higher clonality indicates less diversity [59]. R20 is defined as the fraction of unique clones, in descending order of frequency, that cumulatively account for 20% of the sequenced repertoire: the higher the R20, the less immunodominance there is in a population. A standard quantitative measure of repertoire overlap is Jensen-Shannon divergence (JSD) (2), a tool that accounts for both clone number and frequency and is normalized on a scale of 0 to 1: a JSD of 1 indicates that all clones in 2 populations are distinct; a JSD of 0 indicates that all clones in 2 populations are identical. The code used to analyze TCRβ bulk DNA-seq data and calculate clonality, R20 and JSD is available in previously published paper and has been deposited at https://codeocean.com/capsule/1539294/tree/v2. Clonal overlap of unique sequences among multiple targeted groups was shown in Venn diagrams, which was generated by an online software InteractiVenn (http://www.interactivenn.net) [60]. Cumulative frequency was calculated as a percentage of all sequences weighted by copy numbers in designated populations [61]. Top dominant sequences were ranked by their cumulative frequency within a designated sample.

COVID19-specific hits across genome figure were generated by Adaptive Biotechnologies’ immunoSEQ T-MAP COVID program [62, 63]. Additional statistics and figures were generated using GraphPad Prism (GraphPad Software, La Jolla, CA).

## Data and materials availability

Raw TCRβ bulk DNA-seq data are freely accessible through https://clients.adaptivebiotech.com. The code used to analyze TCRβ bulk DNA-seq data and calculate clonality, R20 and JSD is available in our previously published paper [*Software Impacts*. 2021 (10) 100142] and has been deposited at https://codeocean.com/capsule/1539294/tree/v2.

## Statistical analysis

Graphs were produced, and statistical analyses were performed, using GraphPad Prism version 8.0 (GraphPad Software, Inc., San Diego, CA, USA). Simple linear regression was used to investigate T cell responses of individual antigens. To test the difference in paired observations Wilcoxon matched pairs signed rank test was used, while to compare ranks of unpaired observation Mann-Whitney test was used. P<0.05 is statistically significant.

## Competing interests

J.F serves as a Scientific Consultant for Adaptive Biotechnologies since June 2022. M.P is consultant for Takeda, Clirnet and Rebiotix. A. A receives research funding from Incyte Corporation and consulting fees from Guidepoint Inc and Alphasights.

**Fig. S1:**
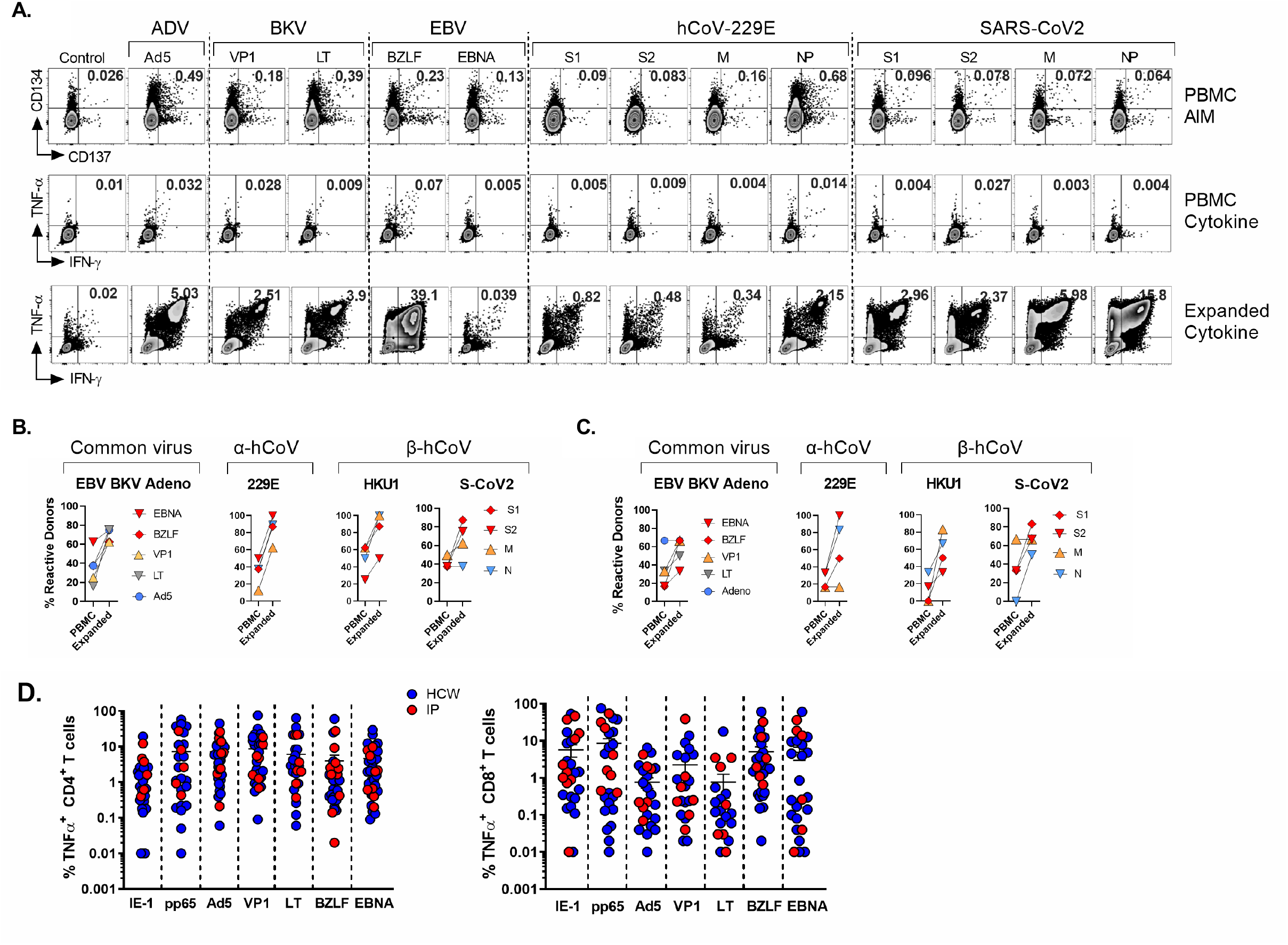
Ex vivo T cell responses as a measure of immunocompetence. PBMCs from each donor (n=8) were stimulated with indicated antigen and frequency of antigen-specific T cells were measured between unexpanded (Day 0) and expanded (Day 14) T cells. **A**. Representative dot plot examples of AIM (day 0, upper row) and cytokine (day 0 and day 14 at middle and lower row, respectively) staining of reactive CD3^+^ T cells. **B**. Comparison of the sensitivity to measure immunocompetence between unexpanded and expanded T cells by ICS assay against indicated antigens (common viruses, α-hCoVs, β-hCoVs & SARS-CoV2). **C**. Comparison of the sensitivity to measure immunocompetence between unexpanded and expanded T cells by AIM assay (CD134^+^CD137^+^CD4^+^ T cells) or ICS assay respectively. **C. D**. Frequencies of virus reactive CD4^+^ T cells (Left panel) and CD8^+^ T cells (Left panel) against indicated antigens in HCW and IP. Blue dots represent HCW, red dots represent IP. Reactive donors were defined as follows: For Day 0 Cytokine release assay and AIM assay, donors displaying more than 0.01% TNF^+^ T cells than the background were considered reactive; while for expanded T cell cytokine assay, donors displaying more than 0.01% TNF^+^ T cells than the background were considered reactive.

**Fig. S2:**
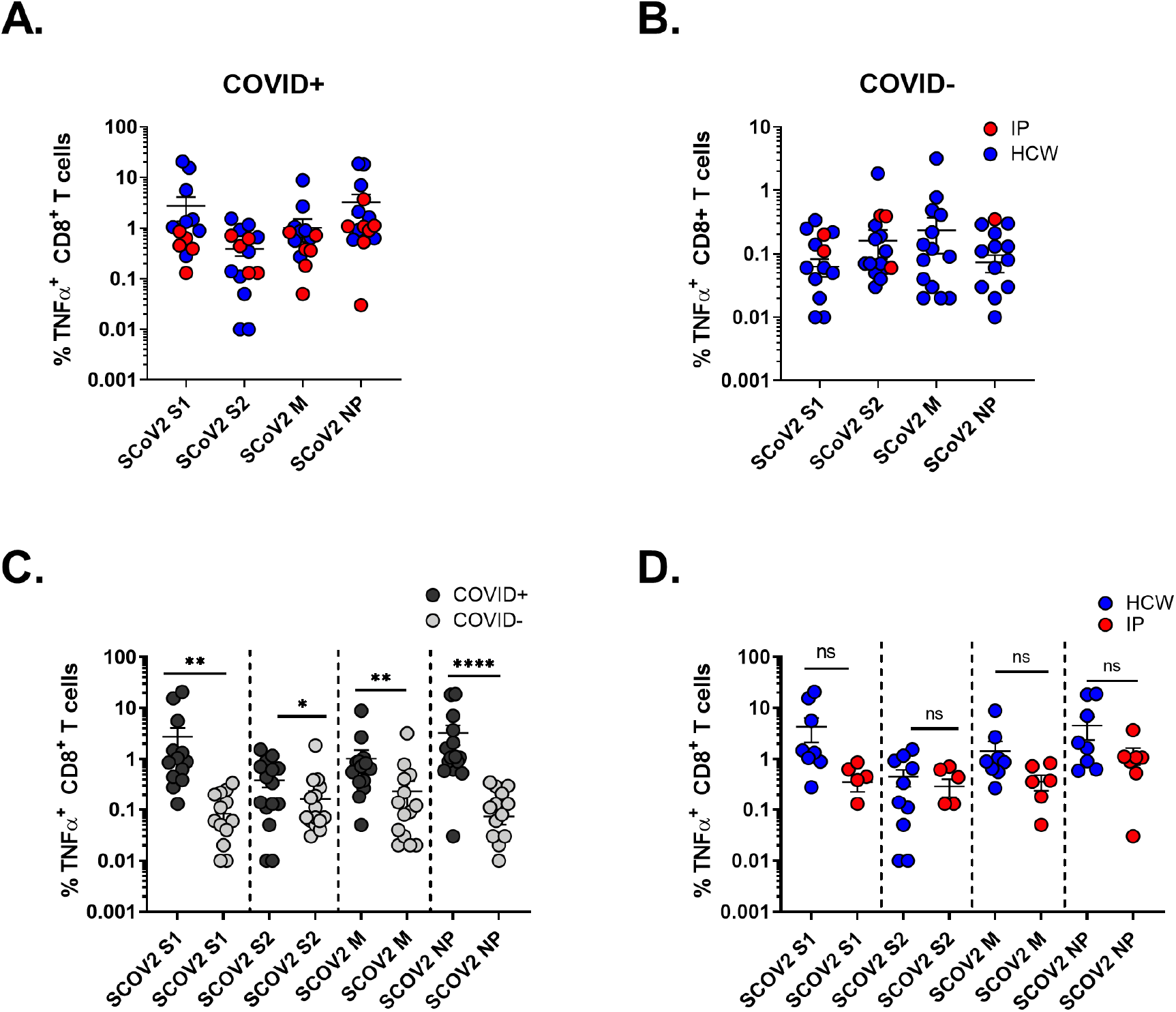
CD8+ T cell responses against SARS-CoV2 antigens. **A**. Frequencies of SARS-CoV2 reactive CD8^+^ T cells against Spike 1, Spike 2, Membrane and Nucleocapsid proteins in COVID+ donors. **B**. Frequencies of SARS-CoV2 reactive CD8^+^ T cells against Spike 1, Spike 2, Membrane and Nucleocapsid proteins in COVID-donors. Blue dots are HCW, red dots are IP. **C**. Comparison of frequencies of SARS-CoV2 reactive CD8+ T cells against indicated proteins between COVID+ and COVID-donors. **D**. Comparison of frequencies of SARS-CoV2 reactive CD8+ T cells against indicated proteins between HCW and IP. Statistically significant differences of reactivity were determined by Mann-Whitney test. **P < 0.01, ***P < 0.001.

**Fig. S3:**
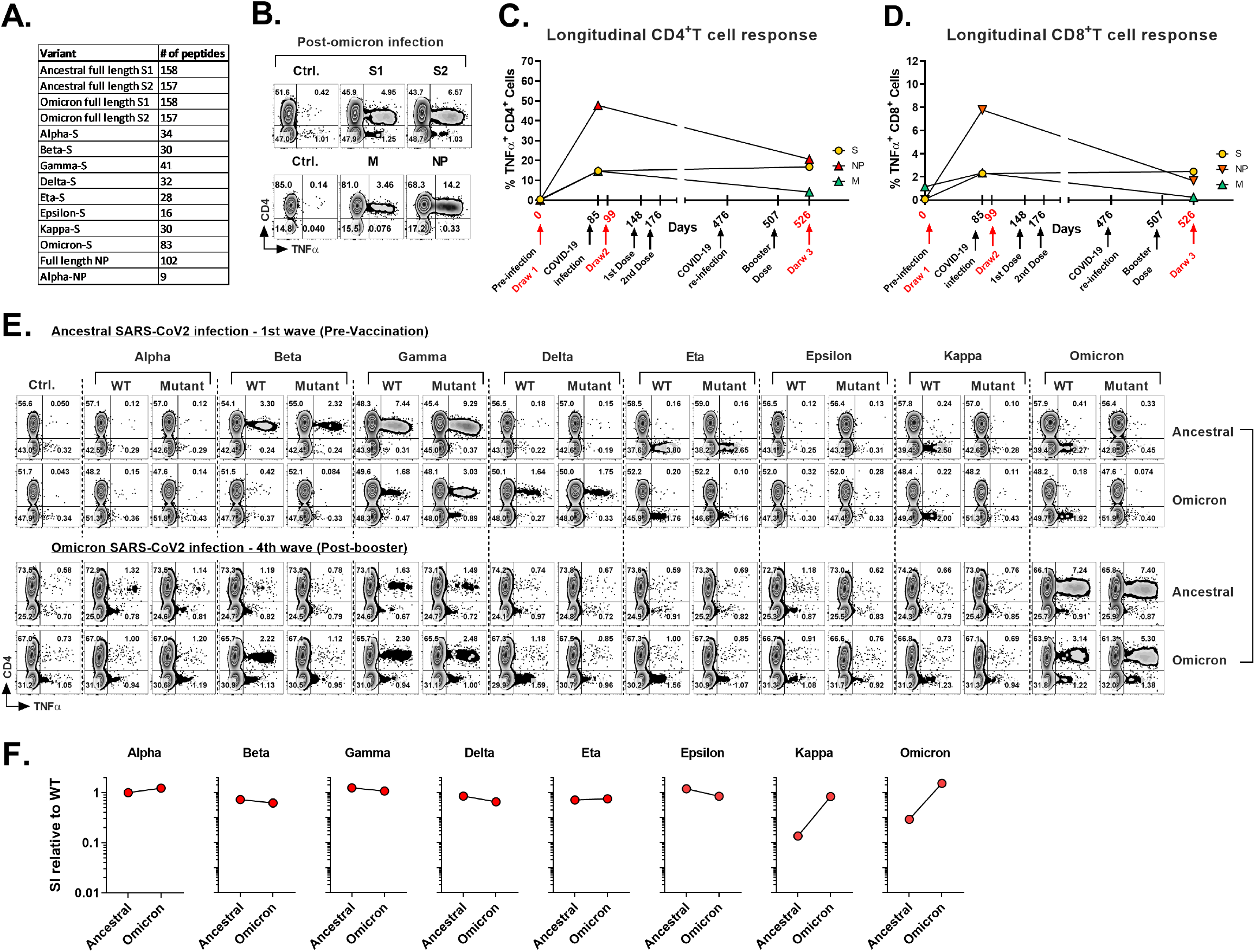
Comparison of T cell response to variants post ancestral and omicron infection in a healthcare worker. **A**. Table showing number of peptides in pepmixes of indicated antigens. **B**. Dot plot showing T cell response against Membrane and Nucleocapsid proteins after omicron infection. **C-D**. Longitudinal analysis of CD4^+^ T cell **(C.)** and CD8^+^ T cell **(D.)** response analysis of donor 1008. **E**. Dot plots depicting T cell response to SARS CoV2 variants. Top panel represents response after infection during 1^st^ wave before vaccination, whereas response after infection during fourth wave is shown in bottom panel **F**. Comparison of T cell response against all variants relative to their WT counterpart after ancestral (top row in top panel) and omicron infection (bottom row in bottom panel).

**Fig. S4:**
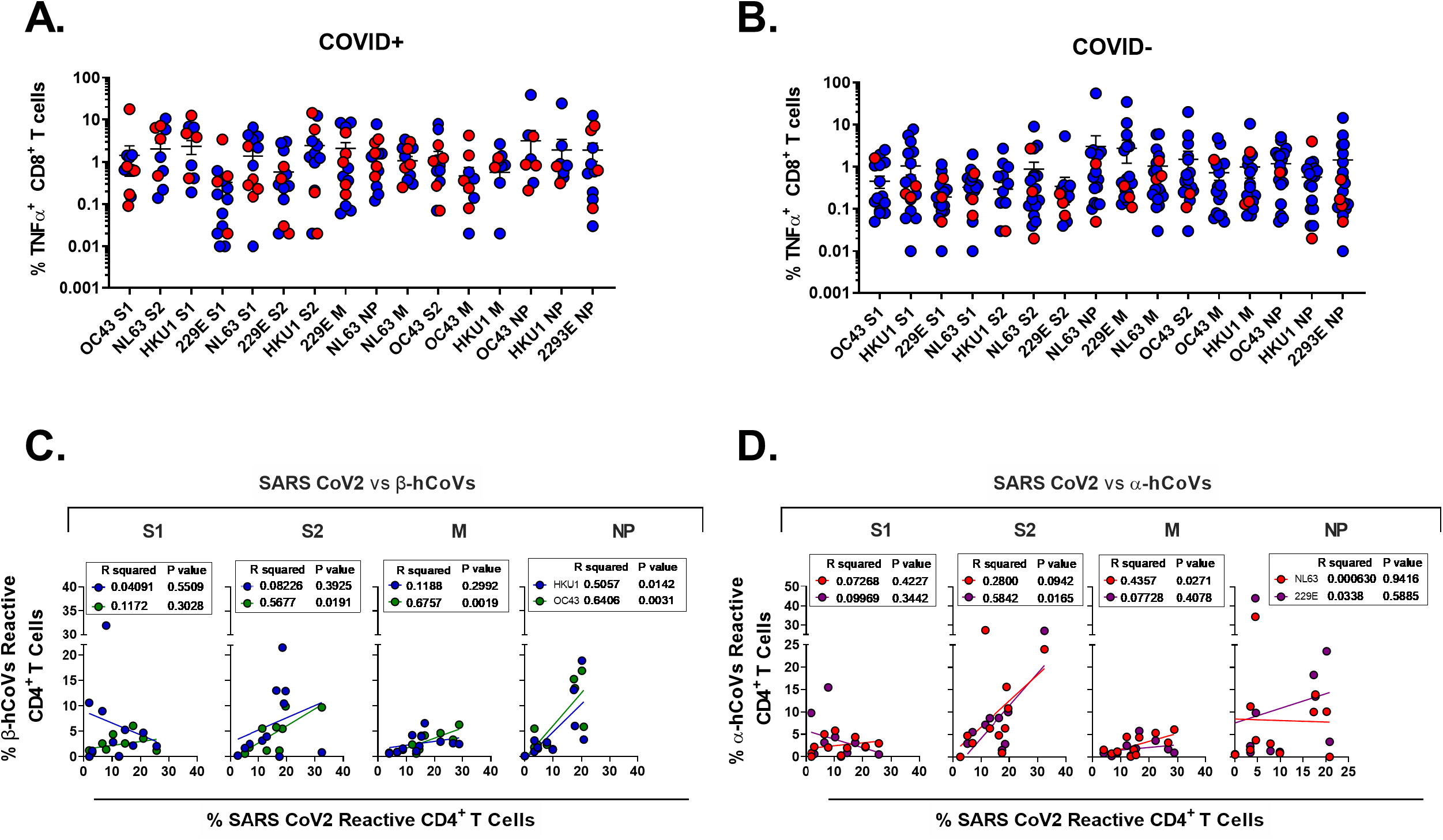
CD8+ T cell responses against SARS-CoV2 antigens. **A**. Frequencies of reactive CD8^+^ T cells against S1, S2, M, and NP proteins from human coronavirus (hCoVS) in COVID+ donors. **B**. Frequencies of reactive CD8^+^ T cells against S1, S2, M, and NP proteins from human coronavirus (hCoVS) in COVID-donors. **C**. Linear correlation between SARS-CoV2 and α-coronavirus reactivity in COVID+ healthcare workers. **D**. Linear correlation between SARS-CoV2 and α-coronavirus reactivity in COVID-healthcare workers.

**Fig. S5:**
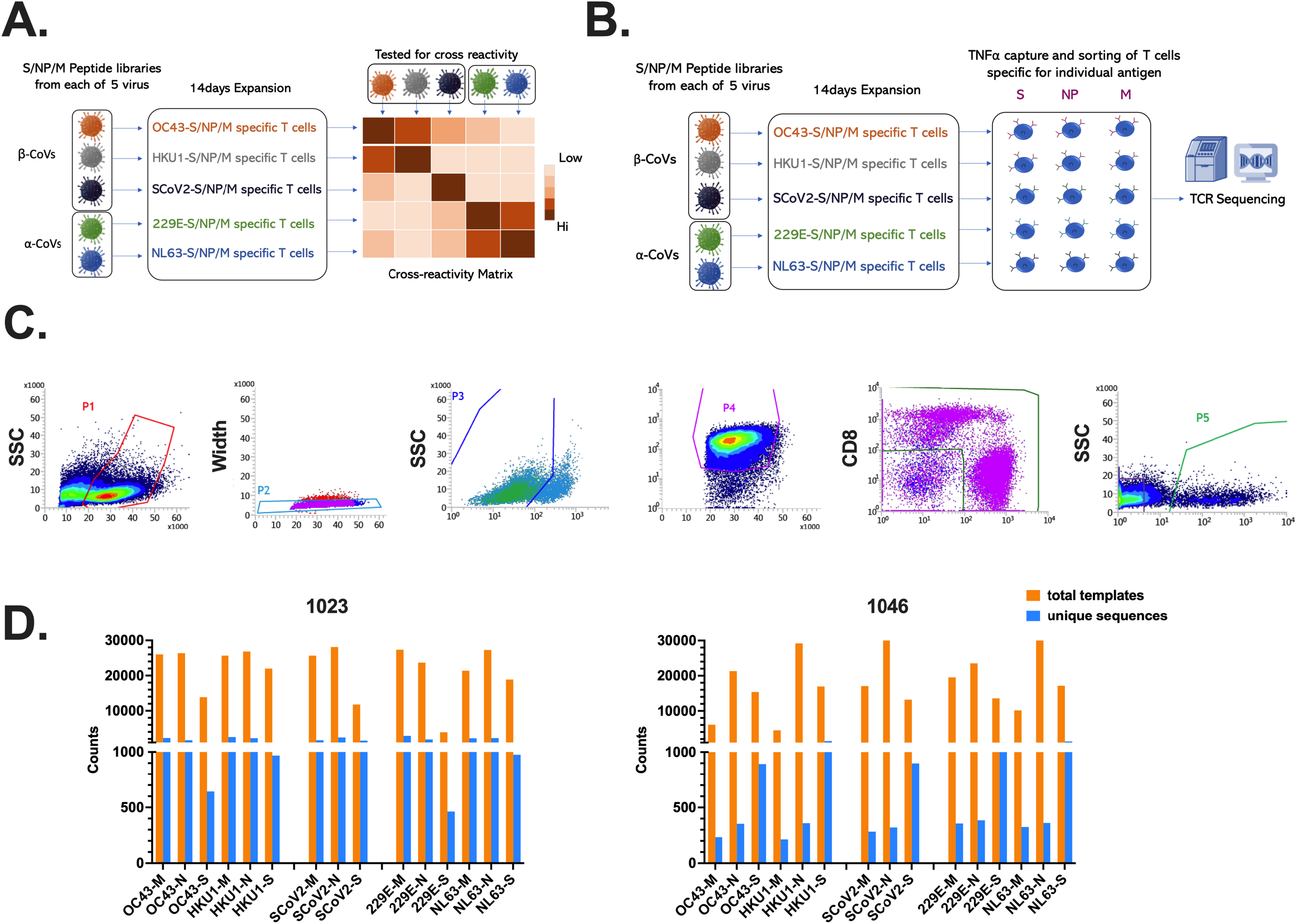
Experimental design for analyzing cross-reactivity and TCR repertoire. **A**. Schematic diagram depicting experimental design to examine cross-reactivity among SARS-CoV2 and human coronaviruses. **B**. Schematic diagram showing strategy for TCR sequencing of antigen-specific T cells. **C**. Gating strategy for sorting virus specific T cells. **D**. Total template counts, and unique sequence counts for TCRs detected in all cultured samples from subject #1023 (Left) and #1046 (Right) against all three antigens (M, N, S) in all 5 CoVs.

**Fig. S6:**
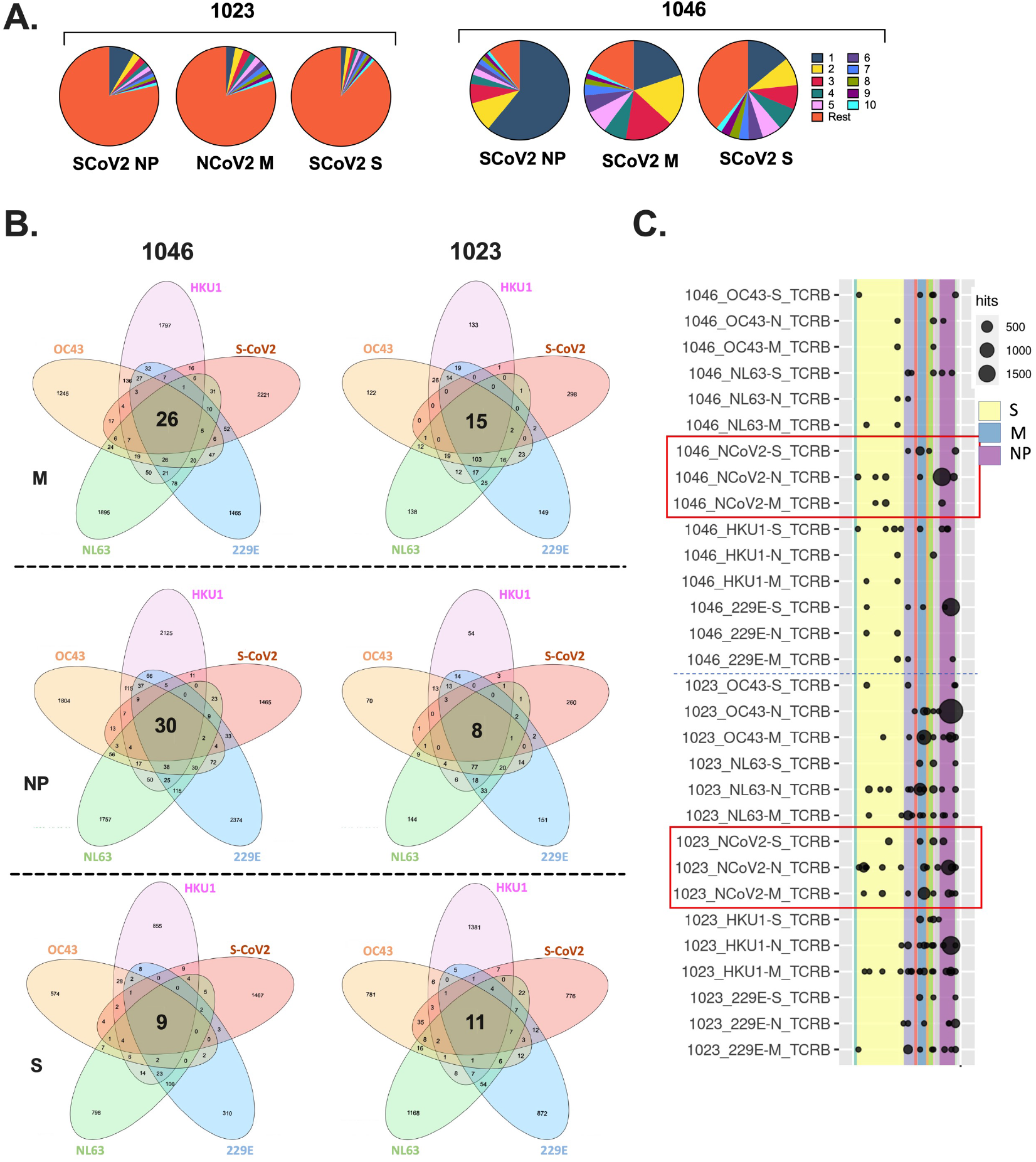
TCR repertoire analysis. **A**. Pie charts showing proportion of top 10 dominant clones within antigen-specific TCR repertoire for both subjects. 1023 (Left) and 1046 (Right). **B**. Venn diagram showing number of TCRs shared between different coronaviruses. **C**. Mapping of Coronavirus specific TCRs in Adaptive Biotechnologies’ ImmuneCODE database containing SARS-CoV-2 specific TCRs identifies shared SARS-CoV2 TCRs along with potential cross-reactive TCRs. **D**.

## References

1. Organization, W.H. WHO Coronavirus (COVID-19) Dashboard. Available from: https://covid19.who.int/.

2. Liu, J., et al., A comparative overview of COVID-19, MERS and SARS: Review article. Int J Surg, 2020. 81: p. 1–8.

3. Breban, R., J. Riou, and A. Fontanet, Interhuman transmissibility of Middle East respiratory syndrome coronavirus: estimation of pandemic risk. Lancet, 2013. 382(9893): p. 694–9.

4. Zumla, A., D.S. Hui, and S. Perlman, Middle East respiratory syndrome. Lancet, 2015. 386(9997): p. 995–1007.

5. Leung, G.M., et al., The Epidemiology of Severe Acute Respiratory Syndrome in the 2003 Hong Kong Epidemic: An Analysis of All 1755 Patients. Annals of Internal Medicine, 2004. 141(9): p. 662–673.

6. Peiris, J.S.M., et al., Clinical progression and viral load in a community outbreak of coronavirus-associated SARS pneumonia: a prospective study. The Lancet, 2003. 361(9371): p. 1767–1772.

7. Gaunt, E.R., et al., Epidemiology and Clinical Presentations of the Four Human Coronaviruses 229E, HKU1, NL63, and OC43 Detected over 3 Years Using a Novel Multiplex Real-Time PCR Method. 2010. 48(8): p. 2940–2947.

8. Corman, V.M., et al., Chapter Eight - Hosts and Sources of Endemic Human Coronaviruses, in Advances in Virus Research, M. Kielian, T.C. Mettenleiter, and M.J. Roossinck, Editors. 2018, Academic Press. p. 163–188.

9. Jo, K.J., et al., Epidemiology and Clinical Characteristics of Human Coronaviruses-Associated Infections in Children: A Multi-Center Study. Front Pediatr, 2022. 10: p. 877759.

10. Masse, S., et al., Epidemiology and Clinical Symptoms Related to Seasonal Coronavirus Identified in Patients with Acute Respiratory Infections Consulting in Primary Care over Six Influenza Seasons (2014-2020) in France. Viruses, 2020. 12(6).

11. Ogimi, C., et al., Prolonged Shedding of Human Coronavirus in Hematopoietic Cell Transplant Recipients: Risk Factors and Viral Genome Evolution. J Infect Dis, 2017. 216(2): p. 203–209.

12. Vardhana, S., et al., Understanding T cell responses to COVID-19 is essential for informing public health strategies. Science immunology, 2022. 7(71): p. eabo1303.

13. Grifoni, A., et al., Targets of T Cell Responses to SARS-CoV-2 Coronavirus in Humans with COVID-19 Disease and Unexposed Individuals. Cell, 2020. 181(7): p. 1489-1501.e15.

14. Tarke, A., et al., Early and Polyantigenic CD4 T Cell Responses Correlate with Mild Disease in Acute COVID-19 Donors. Int J Mol Sci, 2022. 23(13).

15. Rydyznski Moderbacher, C., et al., Antigen-Specific Adaptive Immunity to SARS-CoV-2 in Acute COVID-19 and Associations with Age and Disease Severity. Cell, 2020. 183(4): p. 996-1012.e19.

16. Le Bert, N., et al., Highly functional virus-specific cellular immune response in asymptomatic SARS-CoV-2 infection. Journal of Experimental Medicine, 2021. 218(5).

17. Sekine, T., et al., Robust T Cell Immunity in Convalescent Individuals with Asymptomatic or Mild COVID-19. Cell, 2020. 183(1): p. 158-168.e14.

18. Bange, E.M., et al., CD8+ T cells contribute to survival in patients with COVID-19 and hematologic cancer. Nature Medicine, 2021. 27(7): p. 1280–1289.

19. Jung, J.H., et al., SARS-CoV-2-specific T cell memory is sustained in COVID-19 convalescent patients for 10 months with successful development of stem cell-like memory T cells. Nature Communications, 2021. 12(1): p. 4043.

20. Zhao, J., et al., Airway Memory CD4(+) T Cells Mediate Protective Immunity against Emerging Respiratory Coronaviruses. Immunity, 2016. 44(6): p. 1379–91.

21. Ng, O.-W., et al., Memory T cell responses targeting the SARS coronavirus persist up to 11 years post-infection. Vaccine, 2016. 34(17): p. 2008–2014.

22. Meyerholz, D.K. and S. Perlman, Does common cold coronavirus infection protect against severe SARS-CoV-2 disease? The Journal of Clinical Investigation, 2021. 131(1).

23. Sagar, M., et al., Recent endemic coronavirus infection is associated with less-severe COVID-19. The Journal of Clinical Investigation, 2021. 131(1).

24. Mateus, J., et al., Selective and cross-reactive SARS-CoV-2 T cell epitopes in unexposed humans. Science, 2020. 370(6512): p. 89–94.

25. Kundu, R., et al., Cross-reactive memory T cells associate with protection against SARS-CoV-2 infection in COVID-19 contacts. Nature Communications, 2022. 13(1): p. 80.

26. Araf, Y., et al., Omicron variant of SARS-CoV-2: genomics, transmissibility, and responses to current COVID-19 vaccines. Journal of medical virology, 2022. 94(5): p. 1825–1832.

27. Liu, L., et al., Striking antibody evasion manifested by the Omicron variant of SARS-CoV-2. Nature, 2022. 602(7898): p. 676–681.

28. Grabowski, F., M. Kochańczyk, and T. Lipniacki, The Spread of SARS-CoV-2 Variant Omicron with a Doubling Time of 2.0&ndash;3.3 Days Can Be Explained by Immune Evasion. Viruses, 2022. 14(2): p. 294.

29. Conway, S.R., M.D. Keller, and C.M. Bollard, Cellular therapies for the treatment and prevention of SARS-CoV-2 infection. Blood, 2022. 140(3): p. 208–221.

30. Reiss, S., et al., Comparative analysis of activation induced marker (AIM) assays for sensitive identification of antigen-specific CD4 T cells. PloS one, 2017. 12(10): p. e0186998–e0186998.

31. Dan, J.M., et al., A Cytokine-Independent Approach To Identify Antigen-Specific Human Germinal Center T Follicular Helper Cells and Rare Antigen-Specific CD4<sup>+</sup> T Cells in Blood. The Journal of Immunology, 2016. 197(3): p. 983–993.

32. Bacher, P., et al., Low-Avidity CD4(+) T Cell Responses to SARS-CoV-2 in Unexposed Individuals and Humans with Severe COVID-19. Immunity, 2020. 53(6): p. 1258–1271 e5.

33. Cortese, I., et al., Pembrolizumab Treatment for Progressive Multifocal Leukoencephalopathy. N Engl J Med, 2019. 380(17): p. 1597–1605.

34. Hachmann, N.P., et al., Neutralization Escape by SARS-CoV-2 Omicron Subvariants BA. 2.12. 1, BA. 4, and BA. 5. New England Journal of Medicine, 2022.

35. Tarke, A., et al., SARS-CoV-2 vaccination induces immunological T cell memory able to cross-recognize variants from Alpha to Omicron. Cell, 2022. 185(5): p. 847–859 e11.

36. Reynolds, C.J., et al., Immune boosting by B.1.1.529 (Omicron) depends on previous SARS-CoV-2 exposure. Science, 2022. 377(6603): p. eabq1841.

37. Tarke, A., et al., Comprehensive analysis of T cell immunodominance and immunoprevalence of SARS-CoV-2 epitopes in COVID-19 cases. Cell Rep Med, 2021. 2(2): p. 100204.

38. Soni, M., et al., Development of T-cell immunity in a liver and hematopoietic stem cell transplant recipient following coronavirus disease 2019 infection. Cytotherapy, 2021. 23(11): p. 980–984.

39. Haney, D., et al., Isolation of viable antigen-specific CD8+ T cells based on membrane-bound tumor necrosis factor (TNF)-alpha expression. J Immunol Methods, 2011. 369(1-2): p. 33–41.

40. Migliori, E., M. Chang, and P. Muranski, Restoring antiviral immunity with adoptive transfer of ex-vivo generated T cells. Curr Opin Hematol, 2018. 25(6): p. 486–493.

41. Bollard, C.M., et al., Adoptive immunotherapy for posttransplantation viral infections. Biol Blood Marrow Transplant, 2004. 10(3): p. 143–55.

42. de Candia, P., et al., T Cells: Warriors of SARS-CoV-2 Infection. Trends Immunol, 2021. 42(1): p. 18–30.

43. Liu, R., et al., Decreased T cell populations contribute to the increased severity of COVID-19. Clin Chim Acta, 2020. 508: p. 110–114.

44. Grifoni, A., et al., Targets of T cell responses to SARS-CoV-2 coronavirus in humans with COVID-19 disease and unexposed individuals. Cell. 2020 May 20. doi: 10.1016/j.cell.2020.05.015.

45. Marcus, N., et al., Minor Clinical Impact of COVID-19 Pandemic on Patients With Primary Immunodeficiency in Israel. Frontiers in Immunology, 2021. 11.

46. Yazdanpanah, F., M.R. Hamblin, and N. Rezaei, The immune system and COVID-19: Friend or foe? Life Sci, 2020. 256: p. 117900.

47. Dan, J.M., et al., Immunological memory to SARS-CoV-2 assessed for up to 8 months after infection. Science, 2021.

48. Rucinski, S.L., et al. Seasonality of Coronavirus 229E, HKU1, NL63, and OC43 From 2014 to 2020. in Mayo Clinic Proceedings. 2020. Elsevier.

49. Moss, P., The T cell immune response against SARS-CoV-2. Nature Immunology, 2022. 23(2): p. 186–193.

50. Bacher, P., et al., Low-Avidity CD4+ T Cell Responses to SARS-CoV-2 in Unexposed Individuals and Humans with Severe COVID-19. Immunity, 2020. 53(6): p. 1258-1271.e5.

51. Rock, K.L., E. Reits, and J. Neefjes, Present Yourself! By MHC Class I and MHC Class II Molecules. Trends Immunol, 2016. 37(11): p. 724–737.

52. Nielsen, M., et al., MHC class II epitope predictive algorithms. Immunology, 2010. 130(3): p. 319–28.

53. Kuthuru, O., et al., Rapid induction of antigen-specific CD4 T cells is associated with coordinated humoral and cellular immunity to SARS-CoV-2 mRNA vaccination. Immunity, 2021. 54: p. 2133–2142.

54. Taus, E., et al., Dominant CD8+ T Cell Nucleocapsid Targeting in SARS-CoV-2 Infection and Broad Spike Targeting From Vaccination. Frontiers in immunology, 2022: p. 506.

55. Peng, Y., et al., An immunodominant NP105–113-B*07:02 cytotoxic T cell response controls viral replication and is associated with less severe COVID-19 disease. Nature Immunology, 2022. 23(1): p. 50–61.

56. Jing, L., et al., T cell response to intact SARS-CoV-2 includes coronavirus cross-reactive and variant-specific components. JCI Insight, 2022. 7(6).

57. Woldemeskel, B.A., et al., CD4+ T cells from COVID-19 mRNA vaccine recipients recognize a conserved epitope present in diverse coronaviruses. The Journal of Clinical Investigation, 2022. 132(5).

58. Keller, M.D., et al., SARS-CoV-2–specific T cells are rapidly expanded for therapeutic use and target conserved regions of the membrane protein. Blood, 2020. 136(25): p. 2905–2917.

59. Langerak, A.W., et al., Immunoglobulin/T-cell receptor clonality diagnostics. Expert Opin Med Diagn, 2007. 1(4): p. 451–61.

60. Heberle, H., et al., InteractiVenn: a web-based tool for the analysis of sets through Venn diagrams. BMC Bioinformatics, 2015. 16: p. 169.

61. Fu, J., et al., Lymphohematopoietic graft-versus-host responses promote mixed chimerism in patients receiving intestinal transplantation. J Clin Invest, 2021. 131(8).

62. Snyder, T.M., et al., Magnitude and Dynamics of the T-Cell Response to SARS-CoV-2 Infection at Both Individual and Population Levels. medRxiv, 2020.

63. Nolan, S., et al., A large-scale database of T-cell receptor beta (TCRbeta) sequences and binding associations from natural and synthetic exposure to SARS-CoV-2. Res Sq, 2020.

