## Supplementary File for "The prospect of universal coronavirus immunity: a characterization of reciprocal and non-reciprocal T cell responses against SARS-CoV2 and common human coronaviruses"

**Methods**

**Material Tables:**

**Cytokine Free Medium (CFM):**

| #catalog | Company | Reagent | Concentration |
|---|---|---|---|
| 11-875-119 | Fisher Scientific | RPMI 1640 with L-Glutamine | 47% |
| 12-055-091 | Fisher Scientific | AIM -V | 47% |
| 100-318 | Gemini Bio-Products | Human AB Serum | 5% |
| 10-378-016 | Fisher Scientific | Penicillin Streptomycin Glutamine (100X) | 1% |
| 15-710-072 | Fisher Scientific | Gentamicin (10 mg/mL) | 0.01% |

**Cytokine added to CFM:**

| # Catalog | Company | Name | Final Concentration |
|---|---|---|---|
| 200-07-250UG | Peprotech | IL-7 | 10ng/ml |
| 200-15-250UG | Peprotech | IL-15 | 10ng/ml |
| 170-076-148 | Miltenyi Biotec | IL-2 | 30U/ml |

| **CYTOKINE PANEL** | | | | |
|---|---|---|---|---|
| **#catalog** | **Company** | **Antibody** | **Fluorochrome** | **Concentration** |
| 130-110-206 | Miltenyi Biotec | Viobility 405/520 Fixable Dye | BV510 | 100 |
| 317336 | Biolegend | CD3 | PerCP/Cy5.5 | 100 |
| 100451 | Biolegend | CD4 | PE/Cy7 | 100 |
| 344714 | Biolegend | CD8 | APC/Cy7 | 100 |
| 372222 | Biolegend | Granzyme B | APC | 50 |
| 502909 | Biolegend | TNF-α | PE | 50 |
| 500334 | Biolegend | IL-2 | BV650 | 50 |
| 512326 | Biolegend | IL-17A | BV605 | 50 |
| 502532 | Biolegend | IFN-γ | BV421 | 50 |

| **$CD4^+$ MEMORY PHENOTYPE PANEL** | | | | |
|---|---|---|---|---|
| **#catalog** | **Company** | **Antibody** | **Fluorochrome** | **Concentration** |
| 130-110-205 | Miltenyi Biotec | Viobility 405/452 Fixable Dye | BV421 | 100 |
| 353218 | Biolegend | CCR7 | AF647 | 200 |
| 317336 | Biolegend | CD3 | PerCP/Cy5.5 | 100 |

| | | | | |
|---|---|---|---|---|
| 100451 | Biolegend | CD4 | PE/Cy7 | 100 |
| 344714 | Biolegend | CD8 | APC/Cy7 | 100 |
| 393328 | Biolegend | CD57 | BV711 | 200 |
| 304246 | Biolegend | CD45RO | BV510 | 200 |
| 302808 | Biolegend | CD27 | PE | 200 |
| 305606 | Biolegend | CD95 | FITC | 200 |
| 351334 | Biolegend | CD127 | BV605 | 200 |
| 304136 | Biolegend | CD45RA | BV650 | 200 |
| 304820 | Biolegend | CD62L | AF700 | 200 |
| 329930 | Biolegend | PD-1 | BV786 | 200 |
| 325616 | Biolegend | CD14 | BV421 | 200 |
| 115523 | Biolegend | CD19 | Pacific blue | 200 |

| **$CD4^+$ TH SUBTYPES PANEL** | | | | |
|---|---|---|---|---|
| **#catalog** | **Company** | **Antibody** | **Fluorochrome** | **Concentration** |
| 130-110-206 | Miltenyi Biotec | Viobility 405/520 Fixable Dye | BV510 | 100 |
| 317336 | Biolegend | CD3 | PerCP/Cy5.5 | 100 |
| 100451 | Biolegend | CD4 | PE/Cy7 | 100 |
| 339912 | Biolegend | CD161 | APC | 50 |
| 359412 | Biolegend | CCR4 | PE | 100 |
| 565925 | Biolegend | CCR6 | BV421 | 100 |
| 353722 | Biolegend | CXCR3 | APC/Cy7 | 100 |
| 313713 | Biolegend | CCR5 | AF700 | 100 |
| 310716 | Biolegend | CCR3 | BV605 | 100 |
| 356914 | Biolegend | CXCR5 | FITC | 50 |
